# Class IIa HDACs regulate learning and memory through dynamic experience-dependent repression of transcription

**DOI:** 10.1101/540856

**Authors:** Yongchuan Zhu, Min Huang, Eric Bushong, Sebastien Phan, Marco Uytiepo, Elizabeth Beutter, Daniel Boemer, Kristin Tsui, Mark Ellisman, Anton Maximov

## Abstract

The formation of new memories requires transcription. However, the mechanisms that limit signaling of relevant gene programs in space and time for precision of information coding remain poorly understood. We found that, during learning, the cellular patterns of expression of early response genes (ERGs) are regulated by class IIa HDACs 4 and 5, transcriptional repressors that transiently enter neuronal nuclei from cytoplasm after sensory input. Mice lacking these repressors in the forebrain have abnormally broad experience-dependent expression of ERGs, altered synaptic architecture and function, elevated anxiety, and severely impaired memory. By acutely manipulating the nuclear activity of class IIa HDACs in behaving animals using a chemical-genetic technique, we further demonstrate that rapid induction of transcriptional programs is critical for memory acquisition but these programs may become dispensable when a stable memory is formed. These results provide new insights into the molecular basis of memory storage.

## Introduction

Sensory exprience activates transcriptional programs in the brain that affect neuronal wiring, synaptic function and memory formation ^1,2^. The first waves of dynamic experience-dependent changes in neuronal transcriptomes arise from transient expression of early response genes (ERGs) which are only induced in subsets of cells in a given environment ^3–5^. Most ERGs encode transcription factors (TFs) that exert their physiological effects through downstream target genes ^1^. Labeling for native ERG products and expression of exogenous molecules under the control of ERG promoters is now broadly used in neuroscience to identify and manipulate neuronal ensembles recruited for particular cognitive tasks ^6–8^. For example, memory traces can be artificially retrieved via optogenetic stimulation of ERG-positive cells that were permanently tagged in cortical and hippocampal circuits during learning^9–11^.

While cellular patterns of ERG induction are thought to reflect sparse coding of information in the brain ^3,11,12^, it remains unclear how these patterns are controlled in time and space. Gene expression is orchestrated by TFs and transcriptional repressors that either silence discrete loci or globally alter the state of nuclear chromatin ^13^. Several TFs that mediate activity-dependent synthesis of neuronal mRNAs have been characterized ^1,14,15^, but the extent and molecular mechanisms of negative regulation of this process are poorly understood. It is also unclear how ERGs contribute to different aspects of information processing, including memory acquisition, consolidation, storage and retrieval, since the current knowledge about ERG function in mammals largely stems from experimental paradigms that suffer from poor temporal control ^16–19^.

Class IIa histone deacetylases (HDACs) are a small family of chromatin-binding proteins that shuttle between the nucleus and cytoplasm in a calcium- and phosphorylation-dependent manner ^20–23^. The vertebrate class IIa HDACs interact with class I HDACs, SMRT/NCoR co-repressor complexes and various TFs via their N- and C-terminal domains, and exhibit intrinsically low enzymatic activity on ε-N-acetyl lysine substrates due to evolutionary inactivation of the catalytic site ^24^. Genetic studies have suggested that the class IIa HDAC isoform, HDAC4, is necessary for normal cognition in drosophila, mice, and humans ^25–28^. Yet, prior attempts to define the functional role of HDAC4 in the central nervous system have led to conflicting findings. On the one hand, HDAC4 is capable of binding to TFs essential for synaptic plasticity and memory, CREB, MEF2 and SRF, and its constitutively nuclear gain-of-function mutants repress numerous neuronal genes *in vitro* ^23,25,29^. Puzzlingly, the majority of central neurons retain HDAC4 in cytoplasm ^23,25,30^ and *Hdac4*-deficient mice have been reported to have no detectable abnormalities in mRNA levels in the brain ^26,31^. Despite recent reports that HDAC4 irreversibly enters neuronal nuclei under the pathological conditions, such as Ataxia telangiectasia and Parkinson’s disease ^32,33^, the physiological significance of the nuclear import of this protein in normal circuits is unclear.

Herein, we elucidated the function of class IIa HDACs by combining mouse genetics with imaging, genomic, electrophysiological and behavioral readouts. We found that HDAC4 and its close homolog, HDAC5, redundantly act in the same pathway that is essential for spatiotemporal control of ERG programs during memory encoding. We then developed and applied a new chemical-genetic technique that leverages these findings and permits acute transcriptional repression in brains of behaving animals with a small molecule that rapidly stabilizes HDAC4 in neuronal nuclei. Our results demonstrate that induction of ERGs is necessary for learning, but these genes may not be required after a memory is formed. This work provides insight into the molecular basis of memory storage.

## Results

To explore the possibility that HDAC4 regulates network plasticity during learning we examined the subcellular localization of this protein in wildtype mice after subjecting them to contextual fear conditioning (CFC), a neurobehavioral paradigm that prompts the acquisition of associative fear memory (Fig. 1a,b) ^3,9^. This was accomplished by immunolabeling of brain sections with an antibody whose specificity was validated in *Hdac4* knockouts. Unlike control home-caged mice, trained animals (1.5 hours post-CFC) had a pronounced nuclear accumulation of HDAC4 in 26% to 58% of cells in the somatosensory cortex, areas CA1/CA3 of the hippocampus, and the dentate gyrus, brain regions known to be critical for memory encoding ^2,11,34^. This effect was reversed by 6 hours after CFC, indicating that experience-dependent nuclear import of HDAC4 is transient (Fig. 1c,d and Supplementary Fig. 1).

**Fig. 1.**
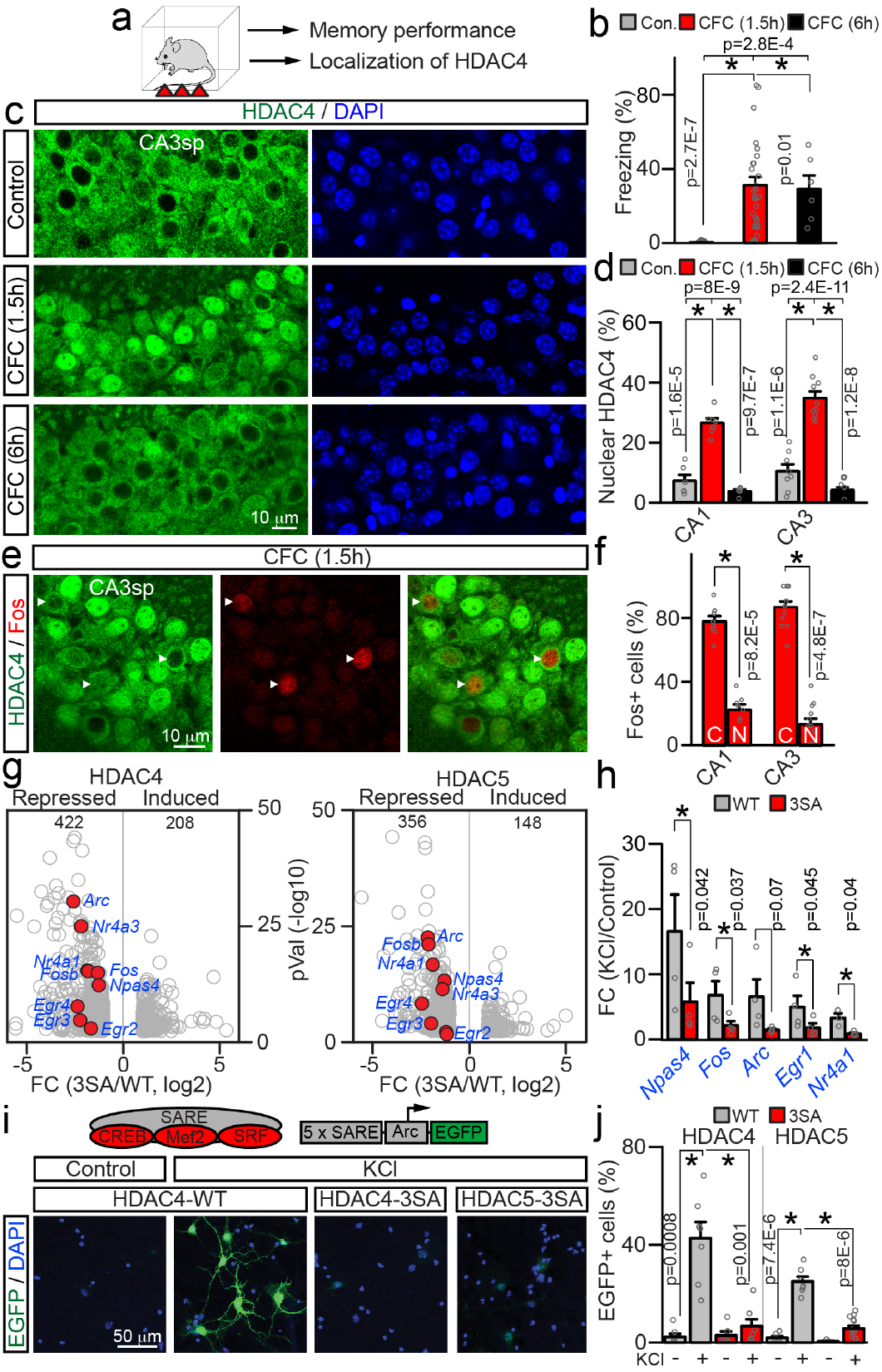
HDAC4 transiently enters neuronal nuclei during associative learning and represses ERGs. **a-f**, Subcellular distribution of native HDAC4 in the hippocampus of wildtype p60 mice before and after contextual fear conditioning (CFC). **a,** Experimental design. **b,** Short-term memory performance was assessed by measuring freezing in the same context. Control, *n* = 11 mice; CFC (1.5h), *n* = 27; CFC (6h), *n* = 6. **c,** Images of HDAC4 immunofluorescence in area CA3sp of DAPI-stained brain sections. See also Supplementary Fig. 1. **d,** Averaged percentages of cells with nuclear HDAC4 in CA1sp and CA3sp. *n* = 4 mice/group (plotted data points are from individual sections). **e,** Brain sections from fear-conditioned animals (1.5h post-CFC) were co-stained for HDAC4 and Fos. **f,** Averaged fractions of Fos-positive cells with cytoplasmic (C) and nuclear (N) HDAC4 in indicated hippocampal subfields (1.5h post-CFC). *n* = 3 mice. **g-j,** Effects of recombinant class IIa HDACs on activity-dependent transcription in primary cortical cultures. Neurons were infected at 2 days *in vitro* (DIV2) with lentiviruses (LVs) that express wildtype HDAC4/5 cDNAs or constitutively nuclear 3SA mutants. **g,** Transcripts were isolated at DIV7 after 30 minute depolarization with KCl (50 mM), and surveyed by deep sequencing. Volcano plots of differentially expressed genes from 2 independent RNA-seq experiments are shown. Fold changes are represented as log2 of 3SA/WT ratio for each HDAC isoform. ERGs are marked in red. See also Supplementary Figs. 2 and 3. **h,** Depolarization-induced expression of ERGs was assessed by quantitative real-time PCR. Averaged FC values from 4 independent experiments are plotted as KCl/Control ratio. **i, j,** Indicated class IIa HDAC LVs were co-infected with the reporter of activity-dependent transcription, E-SARE:EGFP. **i,** Schematics diagram of E-SARE:EGFP and confocal images of EGFP fluorescence in control and KCl-depolarized neurons at DIV7. See also Supplementary Fig. 4. **j,** Summary graphs of reporter induction. *n* = 3 cultures/condition. All quantifications are represented as Mean ± S.E.M. (error bars). * denotes *p* < 0.05 (defined by ANOVA and/or t-test). Actual *p* values are indicated in each graph (vertical for t-test and horizontal for ANOVA).

We then performed double-labeling with antibodies against HDAC4 and Fos to assess the localization of HDAC4 in sparsely distributed Fos-positive neurons that are thought to represent cellular substrates of memory engrams ^3,11^. Remarkably, ∼80% of neurons with detectable Fos immunoreactivity in the hippocampi of fear conditioned mice (1.5 hours post-CFC) retained HDAC4 in cytoplasm, raising the possibility that HDAC4 restricts the expression of Fos and possibly other ERGs in circuits essential for memory storage (Fig. 1e,f).

We and others have previously reported that HDAC4 binds to neuronal chromatin and that prolonged overexpression of a constitutively nuclear phosphorylation-deficient HDAC4 mutant (HDAC4-3SA, Supplementary Fig. 2a) downregulates multiple neuronal genes *in vitro* ^23,25,29^. However, the capacity of HDAC4 for repressing activity-dependent transcription has not been established. We therefore infected cultured cortical neurons with lentiviruses encoding wildtype HDAC4 or HDAC4-3SA and measured the levels of RNAs that were isolated prior to or after brief depolarization with KCl, which stimulates ERGs. Genome-wide mRNA profiling (RNA-seq) and quantitative real-time PCR (qPCR) analyses of individual transcripts consistently demonstrated that HDAC4-3SA interfered with induction of *Fos* and other classical ERGs *Egr2*, *Egr3*, *Egr4*, *Nr4a1*, *Nr4a3*, *Arc*, and *Npas4* ^1,4^ (Fig. 1g,h and Supplementary Figs. 2 and 3).

In addition to HDAC4, neurons in the adult brain abundantly express another class IIa HDAC isoform, HDAC5 ^25^. This isoform has been recently shown to influence reward seeking by interacting with *Npas4* in the Nucleus Accumbens ^35^, but its functional role in other circuits is also unclear. Though we were unable to monitor the dynamic redistribution of native HDAC5 due to insufficient quality of available antibodies, deep sequencing revealed that virtually all ERGs were repressed in cortical cultures by constitutively nuclear HDAC5-3SA mutant as well (Fig. 1g, right panel). Furthermore, the total pools of genes affected by each of the two class IIa HDAC isoforms largely overlapped, raising the possibility that these isoforms are functionally redundant (Supplementary Fig. 2).

To gain a mechanistic insight into how HDACs 4 and 5 silence ERGs, we monitored the expression of EGFP under the control of the *Arc*-based promoter, E-SARE. This short synthetic promoter contains DNA-binding sites for activity-dependent TFs essential for ERG response in the brain, CREB, MEF2, and SRF, and exhibits tight dependence on neuronal excitation *in vitro* and *in vivo* ^36^. Again, KCl-induced expression of E-SARE:EGFP was potently blocked in dissociated cultures by the nuclear 3SA mutants, whereas wildtype cytoplasmic constructs had no effect (Fig. 1i,j and Supplementary Fig. 4). These results imply that class IIa HDACs abolish the ERG transcription through inhibition of TFs on gene promoters.

To determine if HDACs 4 and 5 act in the same pathway in intact circuits, we generated double knockout (DKO) mice that lacked both repressors in glutamatergic neurons throughout the forebrain. These animals carried the previously described *loxP*-flanked (“floxed”) *Hdac4* alleles, constitutive *Hdac5* knockout alleles, and a well-characterized Cre knock-in that drives *loxP* recombination at mid gestation in progenitors of the *Emx1* lineage that give rise to excitatory neurons and astrocytes ^37–40^. (*Hdac4*^*flox/flox*^/*Hdac5*^*−/−*^/*Emx1*^*IRES-Cre*^) (Supplementary Fig. 5a-c). We reasoned that recombination of *Hdac4* in glia should not be problematic since this gene is predominantly expressed in neurons ^25,41^. The homozygote DKOs were born at the expected Mendelian ratio and had regular lifespans. However, they could be easily distinguished from wildtype littermates based on fully penetrant hindlimb clasping in the tail suspension test. In contrast, the clasping was undetectable in mice deficient for individual isoforms (*Hdac4*^*flox/flox*^/*Emx1*^*IRES-Cre*^ and *Hdac5*^*−/−*^), indicating that HDACs 4 and 5 can compensate for each other *in vivo* (Fig. 2a,b).

**Fig. 2.**
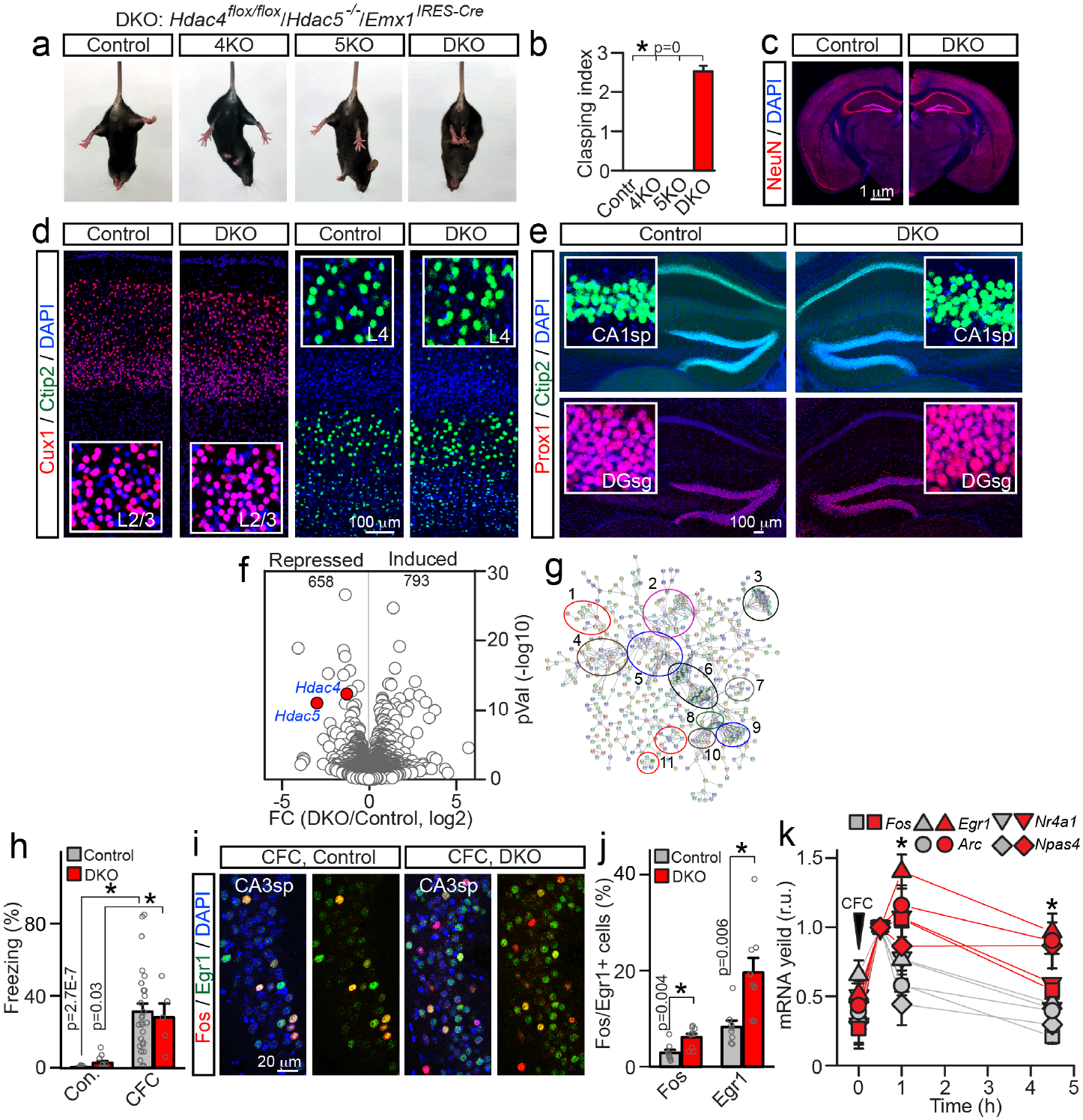
Characterization of HDAC4/5-deficient mice. **a,** Photographs of p60 control, conditional *Hdac4* knockout (KO, *Hdac4*^*flox/flox*^/*Emx1*^*IRES-Cre*^), constitutive *Hdac5* KO (*Hdac5*^*™/−*^) and *Hdac4/5* double knockout (DKO, *Hdac4*^*flox/flox*^/*Hdac5*^*−/−*^/*Emx1*^*IRES-Cre*^) mice subjected to tail suspension test. **b,** Averaged leg clasping indexes. *n* = 20 mice/genotype **c,** Gross brain anatomies of p60 control and DKO mice. Coronal brain sections were stained with DAPI and the antibody to a pan-neuronal marker, NeuN. See also Supplementary Fig. 5. **d, e,** Sections were labeled with antibodies against layer/subtype-specific markers of excitatory neurons, Cux1, Ctip2, and Prox1. Images of the primary somatosensory cortex (**d**) and the hippocampus (**e**) are shown. Higher-magnification frames are displayed in inserts. **f, g,** RNA-seq analysis of gene expression in the hippocampus at p60. Volcano plot (**f**) and the STRING network of differentially expressed transcripts (**g**) are shown (*n* = 3 mice/genotype). The following clusters are marked: 1) Neurotransmitter receptors and calcium channels; 2) Cytoskeleton and scaffolds; 3) Protein turnover; 4) Membrane trafficking; 5) Calcium signaling; 6) G proteins and G protein-coupled receptors; 7) MAP kinases; 8) Adenylate cyclases; 9) Dopamine and hormone receptors; 10) Phosphodiesterases. See also Supplementary Fig. 6. **h-k,** Expression of ERGs in the hippocampus during associative learning. **h,** Analysis of short-term memory performance, assessed by measuring freezing in the same context 1.5h post-CFC. Control/Control, *n* = 11; Control/CFC, *n* = 27; DKO/Control, *n* = 11; DKO/CFC, *n* = 5. **i,** Images of CA3sp of fear-conditioned control and DKO mice (1.5h post-CFC). Sections were labeled with DAPI and antibodies against Fos and Egr1. **j,** Quantifications of Fos/Egr1-positive cells in CA3sp. *n* =3 mice/genotype. **k,** Transcripts were isolated from hippocampi at indicated time-points after CFC, and expression levels of various ERGs were measured by qPCR. Values from 4-5 mice per genotype/timepoint are normalized to 30 min post-CFC to highlight the prolonged window of ERG expression in DKOs. * denotes *p* < 0.05 (t-test). The color coding is the same as in (**j**). All quantifications are represented as Mean ± S.E.M. (error bars). * denotes *p* < 0.05 (defined by ANOVA and/or t-test).

To address the concern that leg clasping of DKO mutants is due to major developmental defects or neurodegeneration, we carried out immunofluorescent imaging of brain sections that were labeled with antibodies for ubiquitous and excitatory neuron subtype-specific markers, NeuN, Cux1, Ctip2, Prox1, and Calbindin. The brains of adult DKOs had undistorted anatomies with appropriate lamination of cortical columns and the hippocampus, and no apparent increase in apoptosis (Fig. 2c-e and Supplementary Fig. 5d). We also examined the cellular architecture of the adult hippocampus by using single cell sequencing (scRNA-seq) on the 10X Genomic platform. Unbiased analysis of 11180 and 12338 cells from dissociated hippocampi of three pairs of control and DKO mice, respectively, showed no changes in the abundance and clustering of neurons, astrocytes, oligodendrocytes, microglia, OPCs, endothelial cells and mural cells, further suggesting that cellular differentiation and survival were unaffected (data not shown).

Next, we investigated the consequences of HDAC4 and HDAC5 loss on gene expression levels by deep sequencing of libraries prepared from pooled hippocampal mRNAs. We chose this approach instead of singe cell sequencing since our scRNA-seq datasets had limited coverage at the expense of large cell numbers. Quantitative analysis of conventional RNA-seq reads with a statistical cutoff at *p* < 0.05 identified 1451 differentially expressed genes, albeit only ∼14% of transcripts were up- or downregulated in class IIa HDAC-deficient animals by more than 2-fold (133 and 75, respectively). As expected, DKOs had a strong depletion of mRNAs of *Hdac4* and *Hdac5* (Fig. 2f). Although the entire pool of affected genes was diverse, the STRING pathway analysis highlighted several prominent clusters of molecules required for synaptic function, network plasticity and memory formation, including various receptors, ion channels and scaffolds, proteins that mediate membrane trafficking and, curiously, multiple adenylate cyclases and phosphodiesterases involved in cAMP signaling (Fig. 2g and Supplementary Fig. 6).

Since the majority of ERGs encode TFs ^1^, these ERGs are repressed by HDACs 4 and 5 *in vitro* (Fig. 1), and each event of transient nuclear import of repressors likely has negligible effect on global expression levels of genes with long-lasting mRNAs, the observed molecular changes in brains of DKO mice may reflect a history of aberrant experience-dependent transcription. To test how loss of HDACs 4 and 5 affects the induction of ERGs, we subjected DKOs to CFC and performed immunolabeling for Fos and Egr1. We imaged area CA3 of the hippocampus where background expression of these genes in home-caged animals is particularly low ^16^. Consistent with observed CFC-induced redistribution of HDAC4 to neuronal nuclei in wildtype animals and analysis of repressor activity in neuronal cultures, DKOs had a ∼2-fold increase in numbers of Fos- and Egr1-positive cells though their contextual freezing responses were comparable to normal littermates (Fig. 2h-j). Likewise, qPCR measurements of transcripts isolated from the whole hippocampi of fear conditioned mice showed that DKOs had increased mRNA levels of *Fos*, *Egr1*, *Nr4a1*, *Arc*, and *Npas4* shortly after CFC, and this increase persisted for several hours (Fig. 2k). These results demonstrate that class IIa HDACs are necessary for spatiotemporal control of ERG programs during memory encoding.

We systematically analyzed the behavior of DKO mutants at p60 to evaluate their motor skills, sensory processing and ability to perform various cognitive tasks. Mice deficient for individual repressor isoforms (*Hdac4*^*flox/flox*^*/Emx1*^*IRES-Cre*^ and *Hdac5*^*−/−*^) were examined in parallel to further define the extent, or lack thereof, of functional redundancy. Despite clasping legs when lifted by tails, DKOs were hardly distinguishable from littermate controls in the standard laboratory environment. However, quantitative measurements of performance in several setups demonstrated that these animals were anxious and they had severely diminished capacity for acquiring new memories. The anxiety-like phenotype was apparent in dark/light transfer tests and analysis of activity in square open fields where DKOs preferred to remain in the dark and avoided the centers of open areas (Fig. 3a-c). These abnormalities were not due to aberrant vision or locomotion, as demonstrated by measurements of head tracks in the optomotor test, total travel distances, and overall horizontal activity (Fig. 3d and Supplementary Fig. 7). When their spatial memory was monitored in the circular Barnes maze ^42^, DKOs were able to identify the escape path during training sessions but showed no target discrimination in the subsequent probe test of memory recall (Fig. 3e-g). In addition, DKOs had virtually no fear extinction, as assessed for a week after CFC on a daily basis by quantifying freezing in the same context (Fig. 3h). The latter also indicates impaired learning rather than inability to erase an existing fear memory ^43^. Notably, none of these phenotypes could be detected in single HDAC4 and HDAC5 KOs (Fig. 3a-c,g,h). It is also noteworthy that the outcomes of spatial and fear memory tests were not merely associated with anxiety and/or aberrant locomotion of DKO mice, as evidenced by quantifications of escape latencies and travel in the Barnes maze and freezing in the irrelevant context (Fig. 3e,f,h; compare freezing in Context A and Context B).

**Fig. 3.**
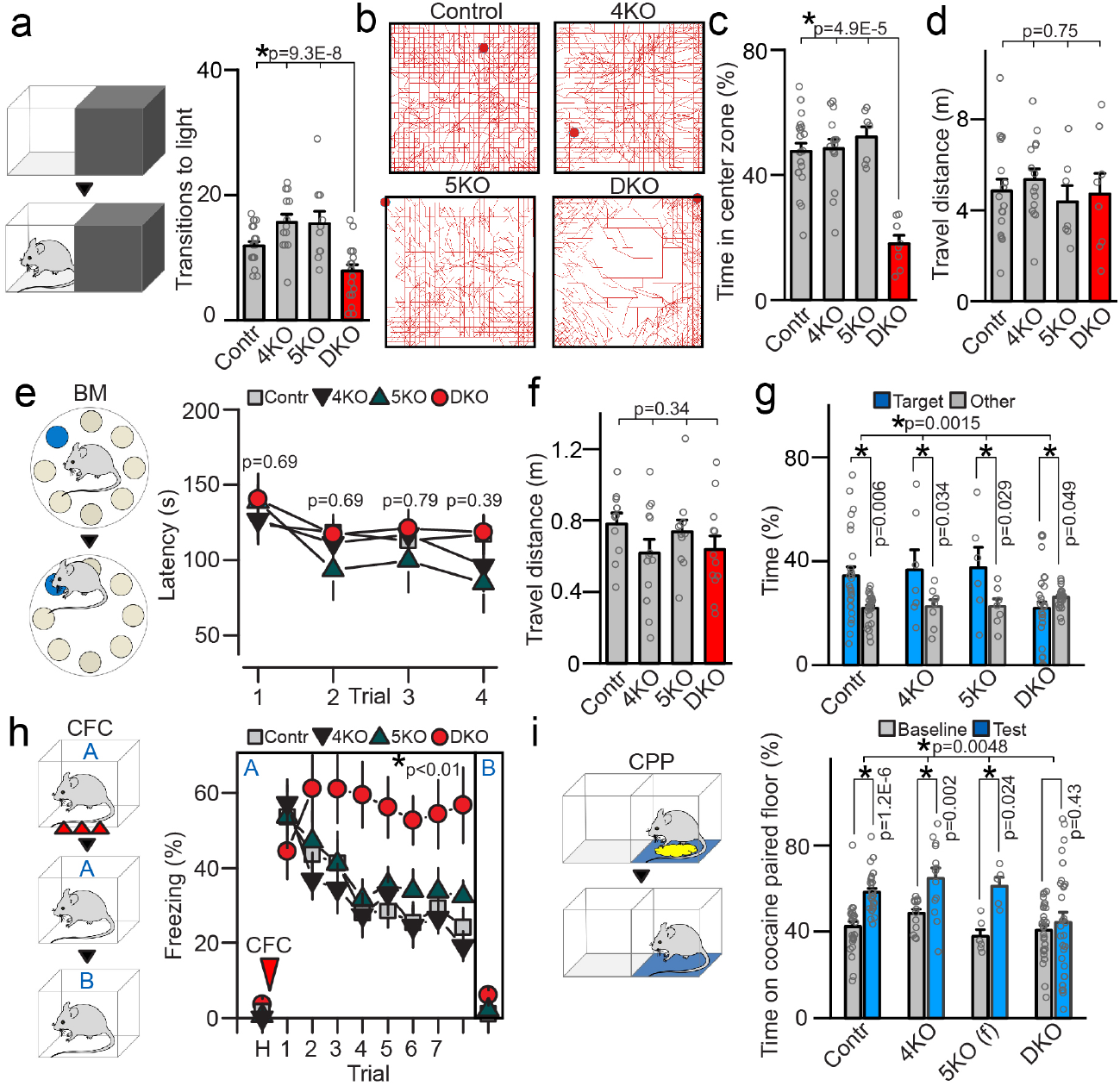
Behavior of class IIa HDAC-deficient mice. Animals of indicated genotypes were examined at p60. See also Supplementary Fig. 7 and methods. **a,** Analysis of anxiety-like behavior in the dark-light transfer test. Schematics of experimental setup and averaged transitions to light are shown. Control: *n* = 19 mice; 4KO: *n* = 13; 5KO: *n* = 10; DKO: *n* = 19. **b-d,** Locomotor activity in the open field setup. **b,** Typical tracks during 30 minute sessions. **c,** Averaged times spent in the center zone. **d,** Total travel distances. Control: *n* = 20; 4KO: *n* = 15; 5KO: *n* = 7; DKO: *n* = 8. **e-g**, Assessment of spatial memory in the Barnes maze. **e,** Schematics of the maze and averaged latencies to escape through the tunnel in each of the four sequential daily training sessions. Control: *n* = 30; 4KO: *n* = 13; 5KO: *n* = 10; DKO: *n* = 30. **f,** Total travel distances in the maze during training. **g,** Memory retrieval in the probe test with escape tunnel removed. Graph shows averaged percentages of time spent in the correct (target) and other quadrants of the maze. Control: *n* = 30; 4KO: *n* = 12; 5KO: *n* = 11; DKO: *n* = 29. **h,** Acquisition and extinction of associative fear memory. Mice were allowed to habituate (H) and then subjected to CFC in Context A. Freezing was measured on a daily basis in the same or irrelevant context (Context B). Control: *n* = 15; 4KO: *n* = 14; 5KO: *n* = 11; DKO: n = 11. **i,** Conditional place preference (CPP) test. Graph shows averaged times spent on cocaine-paired floor. Control: *n* = 27; 4KO: *n* = 13; 5KO: *n* = 6§; DKO: *n* = 29. All quantifications are represented as Mean ± S.E.M. * denotes p < 0.05 (defined by t-test and ANOVA). Actual *p* values are indicated in each graph (vertical for t-test and horizontal for ANOVA). § No sex-related differences in behavior were observed with exception of HDAC5-deficient mice in the CPP test, where only females had preference for cocaine-paired floor.

Because activity of HDAC5 in the Nucleus Accumbens is thought to affect conditional place preference ^35^ we also studied this positive valence behavior in our mouse lines. We found that *HDAC5*^*−/−*^ mutants had no discrimination between regular and cocaine-paired floors, though this defect was only pronounced in males. By contrast, place preference of *Hdac4*^*flox/flox*^*/Emx1*^*IRES-Cre*^ males and females and *Hdac5*^*−/−*^females was similar to that of control animals, but it was completely abolished in DKOs of both genders (Fig. 3i).

To begin to investigate how cognitive deficits of DKO mice correlate with changes in neural circuit structure and function, we analyzed the morphologies of glutamatergic neurons in the hippocampus. We sparsely labeled CA1 pyramidal cells *in vivo* via stereotactic injection of the Cre recombinase-inducible viral tracer, AAV DJ DIO-mGFP (Fig. 4a), collected confocal image stacks from fixed brain sections, and reconstructed single neurons for 3D view. In agreement with our analysis of the anatomy of the hippocampus (Fig. 2e), genetically-tagged HDAC4/5-deficient neurons were appropriately positioned in CA1sp and had fully polarized dendrites (Fig. 4b). Yet, quantitative tracing of dendritic trees revealed significantly higher numbers of branch orders, nodes, ends, and overall complexity of arborization (Fig. 4c). We then imaged dendrites in CA1sr at higher magnification to inspect their spines. While the linear densities of all and mature mushroom-type spines were the same in DKOs and *Emx1*^*IRES-Cre*^ controls, DKO neurons had significantly larger spine sizes (Fig. 4d,e).

**Fig. 4.**
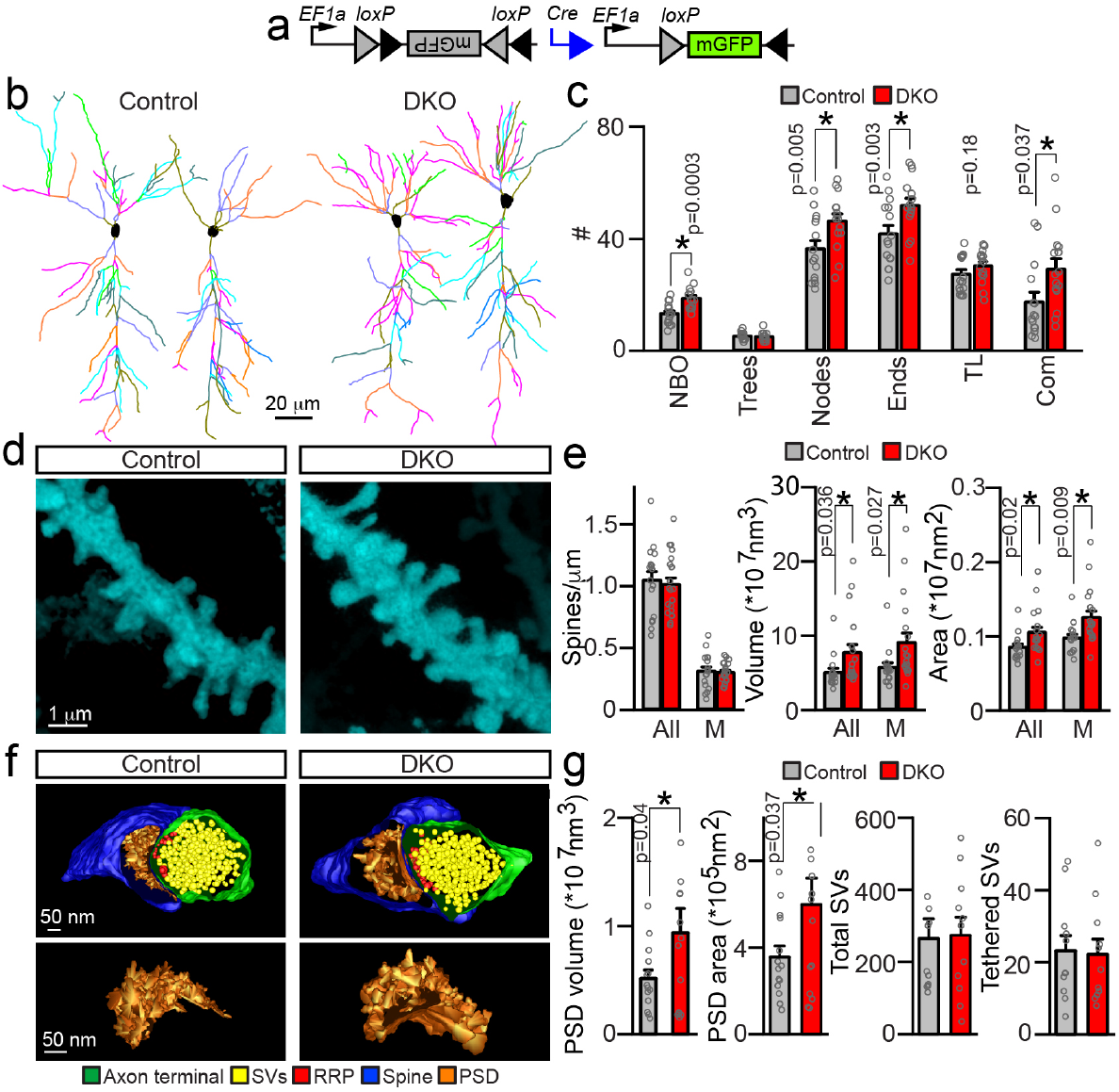
Morphologies of dendrites and synapses of HDAC4/5-deficient excitatory neurons. **a-e,** Pyramidal cells in the CA1 of DKO and control *Emx1*^*IRES-Cre*^ mice were visualized by Cre-inducible expression of membrane-bound GFP from an adeno-associated virus (AAV DJ DIO-mGFP). Animals were injected with AAVs at p45 and analyzed by confocal imaging of brain sections at p60. **a,** Schematic diagram of AAV recombination. **b,** 3D-reconstructions of dendritic trees of individual glutamatergic neurons. **c,** Quantifications of branch orders (NBO), trees, nodes, ends, tree length (TL, μm × 100), and complexity indexes (Com, a.u. x 10000). Averaged values are from 3 mice/15 neurons per genotype. **d,** Pseudo-colored images of postsynaptic spines on proximal dendrites in CA1sr. **e,** Linear densities, surface areas and volumes of all and mushroom (M)-type spines. Control: *n* = 3 mice/17 neurons; DKO: *n* = 3/21. **f, g,** EM tomography analysis of architectures of glutamatergic synapses in CA1sr. **f,** Top panels: 3D reconstructions of spines, postsynaptic densities (PSDs), nerve terminals, and synaptic vesicles (SVs). Bottom panels: Isolated 3D reconstructions of PSDs. **g,** Average volumes and surface areas of PSDs, total SV pools, and numbers of vesicles tethered to active zones (RRP). Control: *n* = 3 mice/14 synapses; DKO: *n* = 3/14. All quantifications are represented as Mean ± S.E.M. * denotes *p* < 0.05 (defined by t-test). Actual *p* values are indicated in each graph.

Considering that spines undergo transient enlargement during long-term synaptic potentiation (LTP) which also coincides with expansion of postsynaptic densities (PSDs) comprised of neurotransmitter receptors, scaffolding molecules and signaling enzymes ^44,45^, we analyzed the ultrastructural organization of glutamatergic synapses in the CA1sr by serial electron microscopy (EM). To this end, tomographic images were acquired from 300 nm sections with 0.5 degree tilt increments at 22,500x magnification, and terminals with opposed spines that contain characteristic PSDs were reconstructed for 3D view from serial stacks. Synapses of HDAC4/5-defienct neurons were appropriately compartmentalized and had unaltered total pools of neurotransmitter vesicles as well as vesicles tethered to presynaptic active zones, but their PSDs were markedly bigger (Fig. 4f,g). Thus, class IIa HDACs restrict dendritic fields of hippocampal pyramidal neurons and affect their spine architectures *in vivo*.

As the next step, we relied on electrophysiological whole-cell recordings from acute hippocampal slices to evaluate intrinsic excitability of CA1 neurons and synaptic transmission in the circuit. Recordings in current clamp mode showed that HDAC4/5-deficient cells had no changes in resting membrane potentials and action potential firing in response to depolarization (Fig. 5a,b). However, measurements of miniature and evoked excitatory postsynaptic currents (EPSCs) demonstrated that loss of the two nuclear repressors leads to augmentation of glutamatergic synaptic strength. Neurons in slices from DKO mice had ∼2-fold higher frequencies of spontaneous miniature AMPA-type EPSCs and elevated AMPA/NMDA ratios of evoked responses (Fig. 5c-f). Together with viral tracing of dendrites of individual neurons, these functional abnormalities indicate both higher numbers of excitatory inputs (due to larger receptive fields) and persistent postsynaptic potentiation since augmented AMPA/NMDA ratio is a signature of LTP ^46,47^. Moreover, we did not detect significant shifts in profiles of synchronous EPSC facilitation/depression during high-frequency stimulus trains, which implies that presynaptic release probability and short-term plasticity of glutamatergic synapses formed by the CA3 afferents were normal (Fig. 5g,h).

**Fig. 5.**
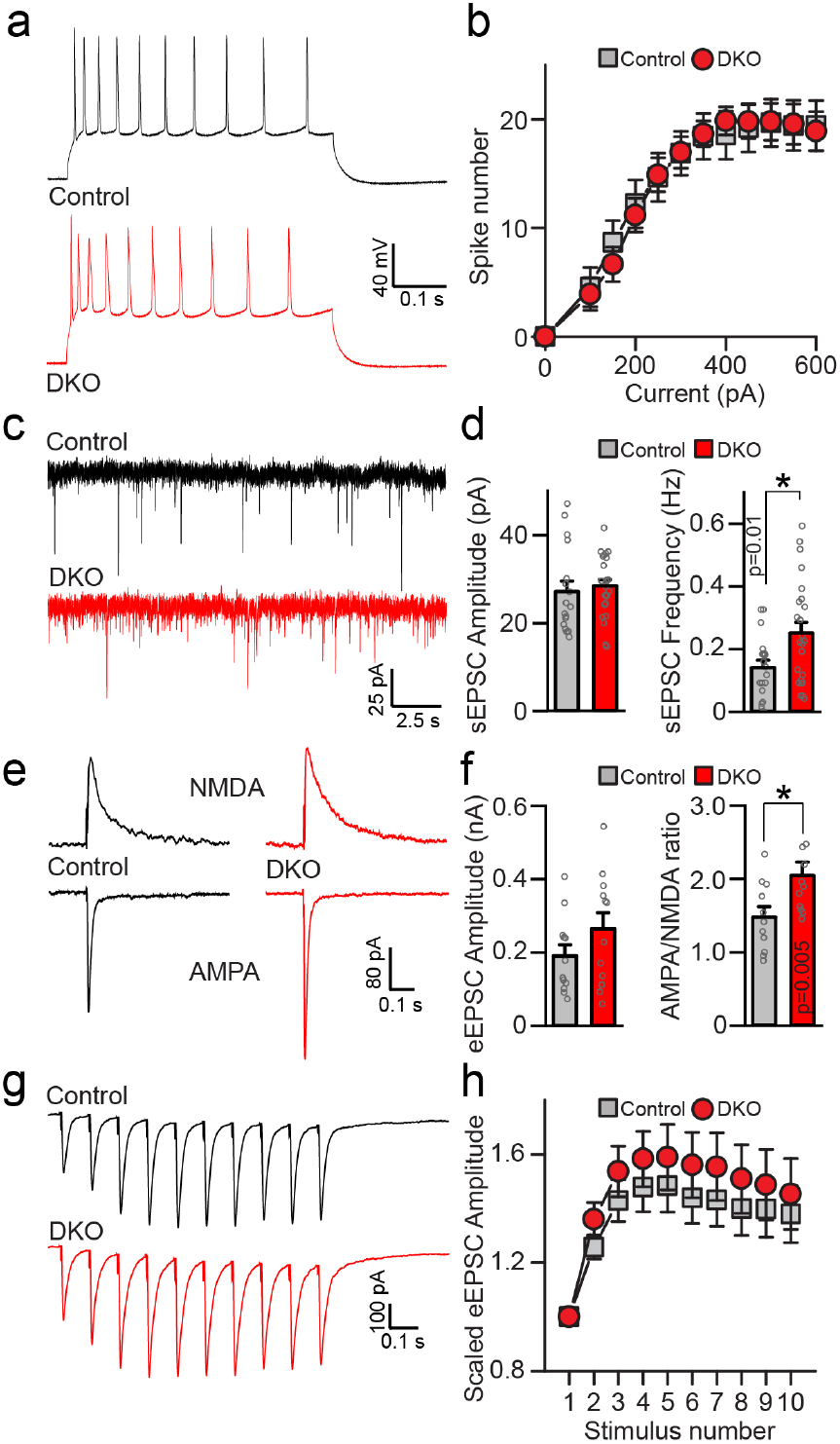
Physiological properties of HDAC4/5-deficient excitatory neurons. Intrinsic excitability and synaptic strength of CA1 pyramidal cells were assessed by electrophysiological whole-cell recordings from acute brain slices from p20 mice. **a,** Sample traces of depolarization-induced action potentials monitored in current-clamp mode. **b,** Averaged numbers of action potentials, plotted as a function of stimulus intensity. Control: *n* = 5 mice/18 neurons; DKO: *n* = 3/27. **c,** Sample traces of spontaneous miniature excitatory postsynaptic currents (sEPSCs) recorded in voltage-clamp mode. Holding potentials were −70 mV. **d,** Averaged sEPSC amplitudes and frequencies. Control: *n* = 5 mice/20 neurons; DKO: *n* = *5*/26. **e,** Traces of evoked AMPA (inward) and NMDA (outward) excitatory postsynaptic currents (eEPSCs) monitored at −70 and +40 mV holding potentials, respectively, in the presence of the GABA receptor blocker, Picrotoxin (50 μM). Synaptic responses were triggered by electrical stimulation of CA3 afferents in the Schaffer collateral path. **f,** Averaged AMPA eEPSC amplitudes (left) and AMPA/NMDA ratios (right). Control: *n* = 3 mice/11 neurons; DKO: *n* = 3/11. **g,** Sample traces of AMPA eEPSC elicited by repetitive stimulation at 10 Hz. **(H)** Averaged eEPSC amplitudes during 10 Hz trains, normalized to amplitudes of first responses. Control: *n* = 5 mice/22 neurons; DKO: *n* = 5/22. All quantifications are represented as Mean ± S.E.M. * denotes *p* < 0.05 (t-test).

Our results thus far provide several important insights into how class IIa HDAC signaling regulates neural circuits at cellular, subcellular and molecular levels, but the contribution of these repressors and their downstream target genes to memory coding is still unclear. Indeed, given the nature of genetic manipulation used in DKO mice, their cognitive defects may be attributed to atypically broad induction of ERGs during learning, pre-existing abnormalities in neuronal wiring, synapse structures and synaptic transmission that reflect repetitive mis-regulation of experience-dependent transcriptional programs and preclude the appropriate processing of sensory information, or both. Likewise, the current understanding of physiological roles of ERGs in the mammalian brain is almost exclusively based on outcomes of prolonged loss- or gain-of-function of these genes in animal models since even drug-inducible Cre drivers and promoters offer relatively poor temporal control ^1,16–18^. While the necessity of gene expression for learning and memory is also demonstrated by pioneering experiments with pharmacological inhibitors of transcription and translation ^2^, these inhibitors lack cellular specificity and globally affect all genes. We therefore devised a strategy for acute transcriptional repression in brains of live and behaving mice that leverages direct stabilization of nuclear HDAC4 with a small molecule.

We fused the constitutively nuclear gain-of-function HDAC4-3SA mutant with the ecDHFR-based N-terminal destabilizing domain (DD), which mediates instant proteasomal degradation of newly synthesized polypeptides ^48^. The decay of this DD-nHDAC4 fusion protein can be blocked by the biologically “inert” antibiotic, Trimethoprim (TMP), that rapidly (within minutes) diffuses through tissues, penetrates the blood-brain barrier, and binds to the DD tag with high affinity ^48,49^ (Fig. 6a). The construct also contained the C-terminal FLAG epitope for detection with antibodies (Supplementary Fig. 8a). To introduce DD-nHDAC4 into genetically-defined neurons, we generated the “floxed” *Rosa26* allele that permits expression under the control of the CAG promoter in a Cre recombinase-dependent manner (*R26*^*floxStop-DD-nHDAC4*^, (Fig. 6a). This allele was then crossed with a well-characterized forebrain-specific driver, *Camk2α*^*Cre* 50^, to induce DD-nHDAC4 in fully differentiated glutamatergic neurons.

**Fig. 6.**
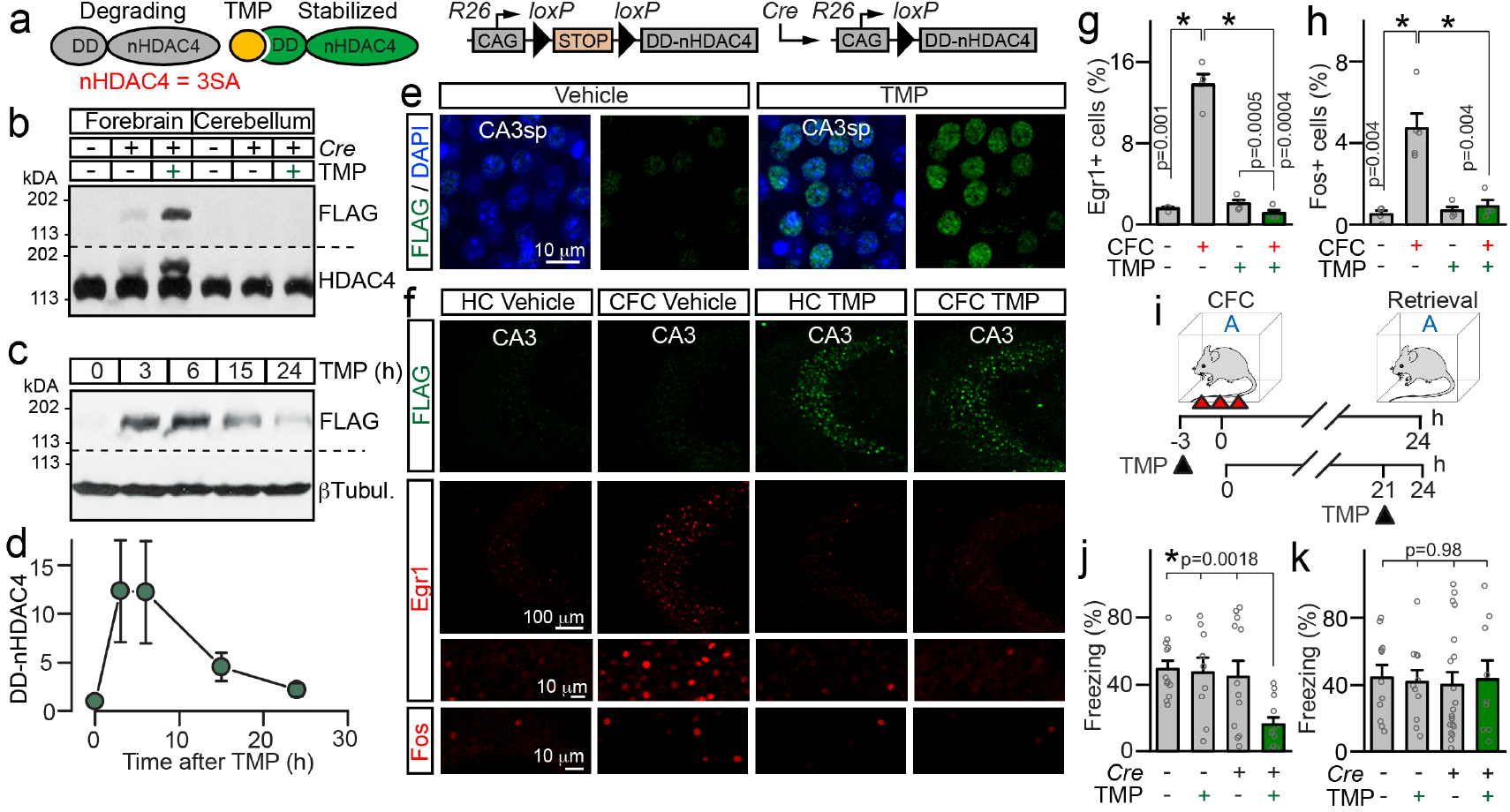
Acute chemical-genetic control of nuclear repression with destabilized HDAC4. **a,** Schematics of stabilization of DD-nHDAC4 protein with TMP and *Rosa26* allele for Cre-dependent expression of DD-nHDAC4 form the CAG promoter (*R26*^*floxStop-DD-nHDAC4*^). See also Supplementary Fig. 8a. **b-k,** DD-nHDAC4 was introduced into excitatory forebrain neurons by crossing *R26*^*floxStop-DD-nHDAC4*^ mice with *Camk2α*^*Cre*^. TMP-lactate (300 μg/gm body weight) or control vehicle solutions were administered via intra-peritoneal injections. **b,** Brains were isolated at p60 3 hours after drug delivery. Cortical and cerebellar protein extracts were probed by immunoblotting with antibodies against FLAG or HDAC4. **c, d,** Time course of DD-nHDAC4 stabilization in the cortex. Immunoblot of samples isolated at different intervals after TMP injection (**c**) and quantifications of DD-nHDAC4 levels (**d**, r.u.) are shown. *n* = 3-4 mice/time point. **e,** Images of DD-nHDAC4 immunofluorescence in CA3sp of vehicle and TMP-treated *R26*^*floxStop-DD-nHDAC4*^/*Camk2α*^*Cre*^ mice (3 hours post-injection). See also Supplementary Fig. 8b. **f-h,** Stabilized DD-nHDAC4 represses ERGs. *R26*^*floxStop-DD-nHDAC4*^/*Camk2α*^*Cre*^ mutants were given single doses of vehicle or TMP. Animals were maintained in home cages (HC) or subjected to CFC (3h post-injection). 1.5h later, brains were sectioned and labeled with antibodies against FLAG, Egr1 or Fos. Panels show images of the CA3 (**f**) and averaged densities of Erg1/Fos-positive cells in this area (**g, h**). See also Supplementary Fig. 8c. **i-k,** Effects of TMP-inducible nuclear repression on associative memory. p60 *R26*^*floxStop-DD-nHDAC4*^/*Camk2α*^*Cre*^ and Cre-negative *R26*^*floxStop-DD-nHDAC4*^ mice were treated with vehicle or TMP prior to CFC or prior to contextual memory retrieval, as depicted in (**i**). **j,** Freezing of animals that were given drugs 3h before learning. Control + vehicle: *n* = 12; Control + TMP: *n* = 9; DD-nHDAC4 + vehicle: *n* = 12; DD-nHDAC4 + TMP: *n* = 12. **k,** Freezing of animals that were given drugs 3h before memory recall. Control + vehicle: *n* = 11; Control + TMP: *n* = 11; DD-nHDAC4 + vehicle: *n* = 19; DD-nHDAC4 + TMP: *n* = 9. All quantifications are represented as Mean ± S.E.M. * denotes *p* < 0.05 (t-test and ANOVA). Actual *p* values are indicated in each graph (vertical for t-test and horizontal for ANOVA).

Immunoblotting and immunofluorescent imaging analyses of brains of adult *R26*^*floxStop-DD-nHDAC4*^/*Camk2α*^*Cre*^ mutants and their control Cre-negative littermates confirmed that DD-nHDAC4 was only expressed in the forebrain in the presence of Cre, was effectively regulated by TMP, and was localized in neuronal nuclei. The protein was nearly undetectable without drug treatments, but its stability was restored within 2-3 hours after single intraperitoneal injections of TMP and was maintained for several hours (Fig. 6b-e). Hence, we were able to achieve temporal control of nuclear class IIa HDAC activity *in vivo* that is at least an order of magnitude faster than temporal control of conventional and inducible promoter-based systems whose kinetics are dictated by rates of mRNA synthesis and degradation. Consistent with the previously described pattern of *Camk2α*^*Cre*^-driven recombination, we observed robust expression of DD-nHDAC4 in the hippocampus, the cerebral cortex, and the amygdala (Supplementary Fig. 8b).

To determine how unstable and TMP-bound DD-nHDAC4 forms affect experience-dependent transcription of class IIa HDAC target genes, we subjected *R26*^*floxStop-DD-nHDAC4*^/*Camk2α*^*Cre*^ mice to CFC and monitored the expression of *Fos* and *Egr1* in the hippocampus by immunofluorescent imaging 1.5 hours later. Analysis of vehicle-treated animals showed that the system has negligible drug-independent background, whereas administration of TMP suppressed the ERG induction in trained mice (Fig. 6f-h and Supplementary Fig. 8c). Hence, this chemical-genetic technique is suitable for transient cell-type-specific silencing of genes that become induced in response to sensory input and, unlike other genes, exhibit narrow peaks of transcriptional activity and encode mRNAs and proteins with short lifespans.

Having validated the new approach, we asked how acute repression of experience-dependent gene programs in glutamatergic neurons impacts associative fear memory using two different protocols in which animals were given TMP either 3 hours before CFC or 3 hours before contextual memory retrieval. In each case, freezing was measured 24 hours after training (Fig. 6i). Both sets of experiments included vehicle- and TMP-treated *R26*^*floxStop-DD-nHDAC4*^/*Camk2α*^*Cre*^ and Cre-negative *R26*^*floxStop-DD-nHDAC4*^ mice to account for potential undesired effects of the antibiotic itself and background activity of TMP-free DD-nHDAC4. Strikingly, transient stabilization of DD-nHDAC4 prior to learning strongly impaired memory, as evidenced by ∼3-fold reduction of freezing times of TMP-injected *R26*^*floxStop-DD-nHDAC4*^/*Camk2α*^*Cre*^ mice (Fig. 6j). This deficit was not due to prolonged baseline leak of the system since vehicle alone had no measurable effect. Moreover, TMP treatments did not affect freezing responses in the absence of *Camk2α*^*Cre*^, excluding the possibility of off-target interactions of the drug (Fig. 6j). Finally, stabilized DD-nHDAC4 had no detectable effect on freezing of pre-conditioned animals that received TMP following CFC, suggesting their memory retrieval was intact (Fig. 6k). Hence, experience-dependent transcriptional programs are required for memory acquisition, but these programs may become dispensable when a stable memory is formed.

## Discussion

In summary, this study elucidates the function of two homologous nuclear repressors, reveals a mechanism that limits experience-dependent expression of genes essential for plasticity of neural circuits, and provides insight into how these genes contribute to memory encoding.

Over the past few years, HDACs have attracted considerable interest as potential therapeutic targets, prompting the design and clinical implementation of small molecule inhibitors ^21,51–53^. However, the precise roles of class IIa HDACs in the nervous system have remained unclear, despite the evidence that loss or mis-localization of these proteins may lead to neurological abnormalities in model organisms and humans ^25–28,30,32,33,35^. Unlike constitutively nuclear class I HDACs, vertebrate class IIa HDACs are unable to deacetylate histones due to evolutionarily acquired Tyrosine to Histidine substitution in the active site of the C-terminal enzymatic domain ^20,24^. The hypothesis that neuronal HDAC4 functions as a transcriptional repressor has been supported by previous findings in several laboratories, including ours, demonstrating that this protein interacts with TFs CREB, MEF2 and SRF and silences multiple genes when artificially forced to neuronal nuclei *in vitro* ^25,29,32,33^. Yet, this hypothesis could not be reconciled with cytoplasmic localization of HDAC4 across the normal brain and perplexing lack of changes in transcription in brains of *Hdac4* knockout mice ^26,30,31,41^. HDAC5 has been recently shown to regulate drug seeking by repressing *Npas4* in the mouse striatum independently of MEF2, though this conclusion was largely based on viral overexpression of nuclear constructs ^35^. To date, the physiological relevance of HDAC5 and mechanism of its action in other circuits were also poorly understood.

In the first part of our work, we showed that HDAC4 is briefly imported to neuronal nuclei during associative learning and that HDAC4 and HDAC5 control the same group of genes, including ERGs. Furthermore, we found that both repressors inhibit a synthetic activity-dependent promoter with binding sites for CREB, MEF2 and SRF, and their simultaneous ablation in excitatory forebrain neurons leads to anxiety and deficits in memory that could not be detected in single KOs. Lastly, we showed that DKO mutants exhibit a broader expression of ERGs after novel experience as well as altered molecular profiles, morphologies and synaptic strength of hippocampal neurons. Taken together, these results suggest that HDACs 4 and 5 redundantly act in the same pathway that restricts the induction of activity-dependent transcriptional programs, thereby limiting the numbers of neurons that undergo structural and/or functional plasticity in response to sensory input. In conjunction with reports that documented aberrant nuclear retention of HDAC4 in cellular and animal models of Parkinson’s disease and Ataxia telangiectasia ^32,33^, our results also imply that class IIa HDACs may exert their effects via different mechanisms under the physiological and pathological conditions depending on kinetics of nuclear import/export. In other words, HDACs 4 and 5 are intrinsically capable of repressing a broad spectrum of targets and they evidently do so upon irreversible nuclear entry (Fig. 1g) ^23,25,33^, but their physiological effects in the brain appear to be strongly biased towards transiently expressed genes with narrow peaks of transcriptional activity and rapidly degrading mRNAs.

A second important aspect of our findings is that they emphasize the critical roles of both acute induction of transcription and spatiotemporal precision of this process for memory formation. Among class IIa HDAC effectors identified in our unbiased genome-wide RNA-seq screens, ERGs *Arc*, *Nr4a1* and *Npas4* are known to be necessary for synaptic plasticity and memory ^16,54–61^. Moreover, the essential roles of CREB, MEF2 and SRF in memory storage have been proven beyond any doubt ^2,14,62–64^. Although the molecular events underlying behavioral phenotypes of repressor-deficient DKO mice are likely complex, it is reasonable to postulate that, at a circuit level, the cognitive deficits of these mice reflect the lack of cellular specificity of information coding that involves long-term changes in wiring and/or synaptic properties of sparsely distributed neural ensembles. On the other hand, our experiments with mice carrying a destabilized TMP-inducible nuclear HDAC4 imply that memory acquisition requires *de novo* transcription within a narrow window after sensory input. From a technical standpoint, these experiments validate a chemical-genetic approach that can be exploited for a broad range of studies in the future. For example, it is feasible to apply targeted expression of destabilized proteins for acute cell type-specific genome editing, induction of TFs, or regulation of neuronal wiring and synaptic strength in live and behaving animals.

While genetic tools leveraging *Fos* and other ERG promoters are becoming increasingly popular for tracing and optogenetic manipulation of neurons with a history of correlated activity elicited by sensory cues ^6–11,65^, recent imaging studies have shown that, in the parietal cortex, activity patterns may also shift over time ^66^. Dynamic spatiotemporal control of ERG programs by class IIa HDACs described herein may contribute to this network flexibility for processing of new information and therefore should be taken into account for better understanding of fundamental principles of neural coding. It is important to note that HDACs 4 and 5 also translocate to neuronal nuclei after global block of NMDA receptors or glutamate release *in vitro* and *in vivo* ^22,23,25,29^, and the circuit-level mechanisms of experience-dependent nuclear import of these repressors need to be further elucidated. It would be of interest to eventually extend the present work with simultaneous *in vivo* imaging of fluorescently tagged class IIa HDACs and calcium indicators, GCaMPs, though this task will require development of new mouse models.

Since class IIa HDACs and ERGs are expressed throughout the brain and in various neuron types, including GABAergic interneurons ^4,25,30,41,60^, it is possible that transcriptional mechanisms described here broadly regulate neural circuits and behavior. It is also noteworthy that vertebrate genomes have two other class II HDAC isoforms whose roles in the nervous system remain unclear: HDAC7, which appears to be predominantly expressed during early development and HDAC9, which encodes a short protein that lacks the C-terminal deacetylase domain but also interacts with MEF2 ^20,25,67^.

## Author Contributions

A.M. and Y.Z. conceived the study. Y.Z. generated mouse lines and personally carried out or supervised all molecular profiling, imaging, and behavioral experiments. M. H. performed electrophysiological recordings from acute brain slices. M. U. and K.T. assisted Y. Z. with confocal microscopy, quantitative real-time PCR, and inhouse tests of associative memory. Other behaviors experiments were performed at the TSRI core facility. E. B. and S.P. prepared samples for EM and collected tomograms. M.E. supervised EM studies. El.B. and D.B. traced genetically-tagged neurons in Neurolucida and analyzed 3D EM reconstructions in IMOD. A.M. wrote the manuscript.

## Acknowledgements

We thank Drs. M. Mayford, L. Stowers and G. Lippi for discussions and critical comments; Drs. E. Olson, R. Shaw and H. Bito for sharing mice and expression vectors; Dr. K. Spencer for assistance with microscopy; Dr. A. Roberts for assistance with behavioral experiments; and The Scripps Research Institute (TSRI) Next Generation Sequencing Core for deep sequencing and bioinformatics support. This study was funded by the NIH grants R01NS087026 (to A.M.), R01GM117049 (to A.M.), R01MH118442 (to A.M.) and P41GM301412 (to M.E.)

## Methods

**Mice** were crossed to produce conditional lines, housed, and analyzed according to protocols approved by the Institutional Animal Care and Use Committee. Strains were maintained in mixed C56BL/6 and 129/Sv backgrounds. Studies were performed with animals of both genders. For each experiment, animal ages are indicated in the main text and/or figure legends. *Ai9, Emx1*^*IRES-Cre*^, *Camk2αCre*, *Hdac4*^*flox/flox*^ and *Hdac5*^*−/−*^alleles were described previously ^37–40,50,68^. To generate the *R26*^*floxStop-DD-nHDAC4*^ allele, the DD-nHDAC4 coding sequence (Supplementary Fig. 8a) was cloned into a targeting vector containing recombination arms, a ubiquitous CAG promoter, a *loxP*-flanked NEO cassette for positive selection, and a Diphtheria toxin cassette for negative selection. DD-nHDAC4 was placed downstream CAG and NEO, which served as a Cre recombinase-removable termination signal (Fig. 6a). The targeting vector was electroporated into ES cells and the positive clones were identified by PCR and Southern Blotting. These clones were then used for blastocyst injection to produce chimeric mice. Chimeras were mated to wildtype mice and their offspring with confirmed germline transmission were used for subsequent crosses with Cre drivers.

### Antibodies and expression vectors

Commercially available antibodies against the following proteins were used for immunofluorescent imaging and immunoblotting. HDAC4 (Santa Cruz, Cat# sc-11418), HDAC4 (Sigma, Cat# H9411), HDAC5 (Sigma, Cat# H4538), βTubulin III (Sigma, T2200), Fos (Santa Cruz, Cat# sc-52G), Egr1 (Cell signaling, Cat# 4153), NeuN (Millipore, Cat# MAB377), Cux1 (Santa Cruz, sc13024), Ctip2 (Abcam, Cat# AB18465), Prox1 (Millipore, Cat# AB5475), Calbindin (Swant, Cat#CB38), Cleaved Caspase3 (Cell Signaling, Cat# 9661). Lentivirus (LV) and Adeno Associated Virus (AAV) shuttle vectors encoding HDAC4-WT, HDAC4-3SA, E-SARE:EGFP and DIO-mGFP were described previously ^25,36,37,50^. To generate LV vectors for expression of WT and 3SA of HDAC5, human coding sequences were amplified by PCR and subcloned downstream of the Synapsin promoter. HDAC5-3SA mutant contained Alanine substitutions of Serine residues S259, S279 and S498.

### Neuronal cultures

Cortices of P1 pups were dissociated by trypsin digestion and seeded onto 24-well plates coated with poly-D-Lysine (Sigma). Cultures were maintained for 4 days in Neurobasal (Invitrogen) supplemented with FBS, B-27 (Invitrogen), glucose, transferrin and Ara-C (Sigma) followed by incubation in the serum-free medium.

### Virus production and infection

Recombinant LVs were produced by co-transfection of human embryonic kidney 293T cells with corresponding shuttle vectors and pVSVg and pCMVΔ8.9 plasmids that encode the elements essential for assembly and function of viral particles. Transfections were performed using the FuGENE reagent (Promega). Secreted viruses were harvested 48 hours later and cleared by brief centrifugation. Cultured neurons were infected at DIV2-5 with 100 μl of viral supernatants per 1 ml of medium. This protocol was optimized to achieve >95% infection efficiency. AAVs were produced in house with shuttle vectors containing EF1α promoter, inverted terminal repeats, the WPRE element, and the hGH polyadenylation signal. To achieve cell type-specific, Cre-inducible expression of fluorescent reporters, coding sequences were flanked by two pairs of loxP sites (DIO) and inserted downstream of a 1.26 kb EF1α promoter in a 3′-5′ orientation. These vectors were packed into AAV serotype DJ using published protocols ^50,69^. Mice were stereotactically injected with 0.5μl of viral stocks via glass micropipettes (10 μm tip diameter) and returned to home cages until experiments.

### Immunohistochemistry

Mice were anesthetized with isoflurane and perfused with 4% PFA. The brains were incubated overnight in 0.5% PFA, and sliced on vibratome in ice-cold PBS. The 90 μm thick, free-floating coronal sections (Bregma −1.4 to −2.5) were briefly boiled in 0.1 M citrate for antigen retrieval. Sections were then washed 3 times in PBS, blocked for 1 hour in a buffer containing 4% BSA, 3% donkey serum, 0.1% Triton, and incubated overnight with primary antibodies diluted in blocking solution, followed by brief washes in PBS and 3 hour incubation with corresponding fluorescently labeled secondary antibodies. Samples were washed again and mounted on glass slides.

### Acquisition and analysis of confocal images

Specific brain regions were annotated using Allen Brain Atlas as the reference (http://mouse.brain-map.org/static/atlas). Images were collected under the Nikon C2 confocal microscope with 10x, 20x and 40x or 60x objectives. Thresholds and laser intensities were established for individual channels and equally applied to entire datasets. Conventional image analysis was conducted with Nikon Elements, FIJI, and Adobe Photoshop software packages. Digital manipulations were equally applied to all pixels. 3D images of dendritic trees of single virally-labeled neurons were collected from 200 μm coronal sections at 0.2 μm Z intervals. Neurons were subsequently reconstructed from serial stacks and analyzed in Neurolucida, as we have previously described ^50^. Spine densities were calculated alone individual fragments of proximal dendrites in CA1sr.

### Electron microscopy

Mice were transcardially perfused with Ringer’s solution followed by perfusion with 150 mM cacodylate, 2.5% glutaraldehyde, 2% paraformaldehyde and 2 mM CaCl_2_. The brains were fixed overnight in the same buffer at 4 °C and cut into 100 μm coronal sections on vibratome. The slices were fixed overnight at 4 °C and then washed for 1 hour in 150 mM cacodylate/2 mM CaCl_2_ on ice. 300 nm thick sections were cut from the SBEM-stained specimens and collected on 50 nm Luxel slot grids (Luxel Corp., Friday Harbor, WA). The grids were coated with 10 nm colloidal gold (Ted Pella, Redding, CA) and imaged at 300 keV on a Titan TEM (FEI, Hillsboro, OR). Double-tilt tilt-series were collected with 0.5 degree tilt increments at 22,500X magnification on a 4k × 4k Gatan Ultrascan camera. Tomograms were generated with an iterative scheme in the TxBR package ^70^. Segmentation of synaptic structures and vesicle pools were performed in IMOD ^70,71^.

### qPCR and deep sequencing

mRNAs were extracted with the RNAeasy Kit (Qiagen). qPCR was performed with the following primers:

*Arc*
Forward: 5’-GGTGAGCTGAAGCCACAAAT-3’
Reverse: 5’-TTCACTGGTATGAATCACTGCTG-3’
*Fos*
Forward: 5’-GGGACAGCCTTTCCTACTACC-3’
Reverse: 5’-AGATCTGCGCAAAAGTCCTG-3’
*Egr1*
Forward: 5’-CCTATGAGCACCTGACCACA-3’
Reverse: 5’-AGCGGCCAGTATAGGTGATG-3’
*Nr4a1*
Forward: 5’-TTGAGTTCGGCAAGCCTACC-3’
Reverse: 5’-GTGTACCCGTCCATGAAGGTG-3’
*Npas4*
Forward: 5’-TTCACAGCTGAGGGGAAGTT-3’
Reverse: 5’-TGTTGAATCGACAACGGAAA-3’
*Gapdh*
Forward: 5’-TCAACGGGAAGCCCATCA-3’
Reverse: 5’-CTCGTGGTTCACACCCATCA-3’

Conventional RNA-seq was performed at the TSRI Next Generation Sequencing Core on the Illumina HiSeq platform. The libraries were generated, barcoded and sequenced according to manufacturer’s recommendations. Sequencing data was analyzed using a two-part in-house pipeline. First, gene expression levels across all samples were quantified with Salmon, a unique quasi-mapping approach for mapping sequences directly to a transcriptome 72. Salmon also includes procedures to remove sample-specific technical biases that can arise during RNA-seq, including sequence-specific, position-specific, and fragment-GC content biases. Next, differential expression analysis was performed in R package DESeq2 ^73^. DESeq2 first adjusts read counts based on a normalization factor that accounts for sample size. This is followed by dispersion estimates based on a negative binomial model which accounts for genes with very few counts. Finally, a Wald test was performed to calculate differential expression between the sample groups. Genes with an adjusted *p* value of < 0.05 and absolute fold change > 2 were identified as significantly differentially expressed.

For single-cell RNA-seq, the brains were placed in oxygenated artificial cerebrospinal fluid (ACSF) containing 125 mM NaCl, 2.5 mM KCl, 2 mM NaH_2_PO_4_, 25 mM NaHCO_3_, 1.3 mM MgCl_2_, 2 mM CaCl_2_, 10 mM Glucose, pH 7.4 (95% O2 and 5% CO2). 300 μm sections were cut on a vibratome (Leica VT1200S) and transferred into ice-cold Hibernate A/B27 medium (50 ml Hibernate A medium with 1 ml B27 and 0.125 ml Glutamax) containing 5 μg/ml actinomycin D (Sigma). Hippocampi were microdissected and incubated for 20 min on ice in Hibernate A/B27 medium containing 1 μM TTX (Tocris), 100 μM AP-V (Tocris) and 5 μg actinomycin D (Sigma). Enzymatic digestion was performed in Hibernate A-Ca medium with 2 mg/ml papain, 1 μM TTX, 100 μM AP-V, 25 μg actinomycin D at 30°C for 40 min. The tissues were then dissociated by gentle trituration through Pasteur pipettes with fire-polished tips. Single-cell suspensions were loaded on a 4-layer OptiPrep gradient and centrifuged at 800 g for 15 min at 4°C. Cell debris were removed by Debris Removal Solution (Miltenyi Biotec) following manufacturer’s instruction. Chromium scRNA-Seq reads were analyzed using Cell Ranger, a suite consisting of multiple pipelines for end to end analysis of single cell data (https://support.10xgenomics.com). In brief, it uses a custom built wrapper around Illumina’s bcl2fastq (https://support.illumina.com) to demultiplex raw basecalls. This is followed by removal of duplicates using UMI counting. These preprocessed samples are aligned using STAR ^74^, which performs splicing-aware alignment of reads to the genome. Aligned reads are then bucketed into exonic and intronic regions using a reference GTF file. Finally, read counts per gene are calculated, and these values are used for downstream estimation of differentially expressed genes.

### Electrophysiology

Mice were anesthetized with isoflurane. Brains were removed and placed into ice-cold oxygenated buffer (95%O_2_/5%CO_2_) containing 110 mM Sucrose, 87 mM NaCl, 2.5 mM KCl, 0.5 mM CaCl_2_, 7 mM MgCl_2_, 25 mM NaHCO_3_, 1.25 mM NaH_2_PO_4_, and 20 mM Glucose. Transverse, 350 μm thick slices were cut with a vibratome and then allowed to recover for ∼1 hour in oxygenated ACSF at 24°C prior to recording. Membrane potentials and synaptic currents were monitored in whole-cell mode using a Multiclamp 700B amplifier (Molecular Devices, Inc.). Recordings were performed at room temperature. For voltage-clamp experiments, the pipette solution contained 122.5 mM C_6_H_12_O_7_, 122.5 mM CsOH, 10 mM CsCl, 1.5 mM MgCl_2_, 5 mM NaCl, 1 mM EGTA, 5 mM HEPES, 3 mM MgATP, 0.3 mM NaGTP, 5 mM QX-314, and 10 mM Na_2_phosphocreatine (pH 7.4). For current-clamp experiments, the pipette solution contained 120 mM K-Gluconate, 20 mM KCl, 4 mM Na_2_ATP, 0.3 mM Na_2_GTP, 4mM Na_2_phosphocreatine, 0.1 mM EGTA, and 10 mM HEPES (pH 7.4). Synaptic release was triggered by 1 ms current injections through the local extracellular stimulating electrode (FHC, Inc. CBAEC75). The frequency, duration, and magnitude of extracellular stimuli were controlled by Model 2100 Isolated Pulse Stimulator (A-M Systems, Inc.). Traces were analyzed offline with pClamp10 (Molecular Devices, Inc.) and Origin8 (Origin Lab) software packages.

### Drug delivery

TMP-lactate (Sigma, Cat #T0667) was reconstituted in PBS prior to each experiment (30 mg/ml) and administered to mice by intraperitoneal injections through a 29g needle at a dose of 300 μg/gm body weight.

### Behavior

<u>Hindlimb clasping</u>. Mice were grasped by tails and lifted for 10 seconds. The scores were assigned using the following criteria, essentially as described^75^: 0 = hindlimbs are consistently splayed outward, away from the abdomen; 1 = one hindlimb is retracted toward the abdomen for more than 50% of the time suspended; 2 = both hindlimbs are partially retracted toward the abdomen for more than 50% of the time suspended; 3 = hindlimbs are entirely retracted and touching the abdomen for more than 50% of the time suspended.

<u>Locomotor activity</u> was measured for 2 hours in polycarbonate cages (42 × 22 × 20 cm) placed into frames (25.5 × 47 cm) mounted with two levels of photocell beams at 2 and 7 cm above the bottom of the cage (San Diego Instruments, San Diego, CA). These two sets of beams allow for the recording of both horizontal (locomotion) and vertical (rearing) behavior.

<u>Vision</u> was assessed in a stationary elevated platform surrounded by a drum with black and white striped walls. Mice were placed on the platform to habituate for 1 minute and then the drum was rotated at 20 rpm in one direction for 1 minute, stopped for 30 seconds, and then rotated in the other direction for 1 minute. The total number of head tracks (15 degree movements at speed of drum) was recorded. In our hands, mice that have intact vision track 5-25 times, whereas blind mice do not track at all.

<u>Anxiety-like behavior</u> was assessed using a light/dark transfer test, which capitalizes on the conflict between exploration of a novel environment and the avoidance of a brightly lit open field ^76^. Mice were placed in the rectangular box divided by a partition into two environments. One compartment (14.5 × 27 × 26.5 cm) was dark (8-16 lux) and the other compartment (28.5 × 27 × 26.5 cm) was highly illuminated (400-600 lux) by a 60 W light source located above it. The compartments were connected by an opening (7.5 × 7.5 cm) located at floor level in the center of the partition. The time spent in the light compartment and the number of transitions into the light compartment during 5 minute sessions were measured as predictors of anxiety.

<u>Spatial memory</u> was examined in the Barnes maze ^42,77^, a setup that consists of an opaque Plexiglass disc 75 cm in diameter elevated 58 cm above the floor by a tripod. Twenty holes, 5 cm in diameter, are located 5 cm from the perimeter, and a black Plexiglass escape box (19 × 8 × 7 cm) is placed under one of the holes. Distinct spatial cues are located around the maze and are kept constant throughout the study. Acquisition: On the first day of testing, a training session was performed, which consisted of placing the animals in the escape box for 1 minute. At the beginning of each session, mice were then placed in the middle of the maze in a 10 cm high cylindrical black start chamber. After 10 seconds the start chamber was removed, a buzzer (80 dB) and a light (400 lux) were turned on, and mice were set free to explore the maze. The session ended when mice entered the escape tunnel or after 3 minutes elapsed. When mice entered the escape tunnel, the buzzer was turned off and animals were allowed to remain in the dark for 1 minute. If a mouse did not enter the tunnel by itself, it was gently put in the escape box for one minute. The tunnel was always located underneath the same hole (stable within the spatial environment), which was randomly determined for each mouse. Mice were tested once a day for 4 days for the acquisition portion of the study. <u>Probe test</u>: For the 5^th^ test (probe test), the escape tunnel was removed and mice were allowed to freely explore the maze for 3 minutes. The time spent in each quadrant was determined and the percent time spent in the target quadrant (the one originally containing the escape box) was compared with the average percent time in the other three quadrants. This is a direct test of spatial memory as there is no potential for local cues to be used in the mouse’s behavioral decision.

<u>Associative memory</u> was examined by using contextual fear conditioning ^3,9,10,78^. Mice were allowed to explore the fear conditioning boxes (Context A, Med Associates SD) for 3 minutes and were then subjected to 4 bursts of foot shocks (0.55 mA, 1 minute inter-shock intervals). Memory tests consisted of one 3 minute exposure to the training box or irrelevant box (Context B). Freezing was determined in 0.75 second bouts and expressed as percent time in the context.

<u>Conditional place preference</u> ^79^ test involves pairing a distinct environmental context (floor type) with a motivationally significant event (cocaine injection). Rectangular Plexiglas black matte boxes (42 × 22 × 30 cm) divided by central partitions into two chambers of equal size (22 × 22 × 30 cm) were used. Distinctive tactile stimuli were provided in the two compartments of the apparatus. One chamber had no additional flooring and the other had lightly textured milky-colored flooring. During pre-conditioning and testing sessions, an aperture (4 × 4 cm) in the central partition allowed the animals to enter both sides of the apparatus. All testing occurred during the dark cycle under red light and was analyzed from video files using Noldus Ethovision software. The experiment consisted of three phases: pre-conditioning, conditioning and testing in the following sequence. Day 1: Pre-conditioning phase with access to both compartments, Day 2-7: Conditioning phase - drug or saline administration followed by immediate confinement in one compartment of the place conditioning apparatus, and Day 8: Test with access to both compartments in the drug-free state. For the pre-conditioning phase, each animal was placed in one compartment of the apparatus and allowed to explore freely the entire apparatus for 30 min. The time spent in each of the two compartments was measured. Mice showing unbiased exploration of the 2 sides of the apparatus (between 45 and 55% time spent on each side) were randomly assigned a chamber in which to receive cocaine. Mice showing biased exploration of either side were given cocaine in the least preferred compartment. Bias was observed in half of the mice tested and this was spread equally across each genotype and sex and therefore did not confound the results. On the following 6 days, 30 min conditioning sessions were given in which animals were injected i.p. with either saline or 10 mg/kg cocaine and immediately confined to one side of the apparatus (alternating these treatments and sides each day). In this way, each mouse experienced 3 pairings of cocaine with one of the apparatus sides. On the day after the final conditioning trial, each mouse was allowed to explore the entire apparatus in a non-drugged state for 30 min and the time spent in each of the two compartments of the conditioned place preference apparatus was recorded. The percent time spent in cocaine side was compared with the pre-conditioning session to examine the development of preference.

### Quantifications and statistical analysis

Means and standard errors were calculated in Origin8. *p* values were determined with Student’s *t*-test (for two groups) and ANOVA (for multiple groups). Quantifications of gene expression levels, dendrite morphologies, spine densities and shapes, synapse ultra-structures and all behavioral experiments were performed in a “blind” manner by investigators who were unaware of genotypes. Details, including sample sizes, can be found in figure legends.

## Notes

#### Summary of Updates

list of authors, addition of new data, correction of statistics, correction of several minor typos.

