## Supplementary material for "Class IIa HDACs regulate learning and memory through dynamic experience-dependent repression of transcription": Combined supplementary matherials w/o raw gene expression data (from another version of this preprint: version 1, 2019-02-05)

**Supplementary materials** for Zhu el al.,

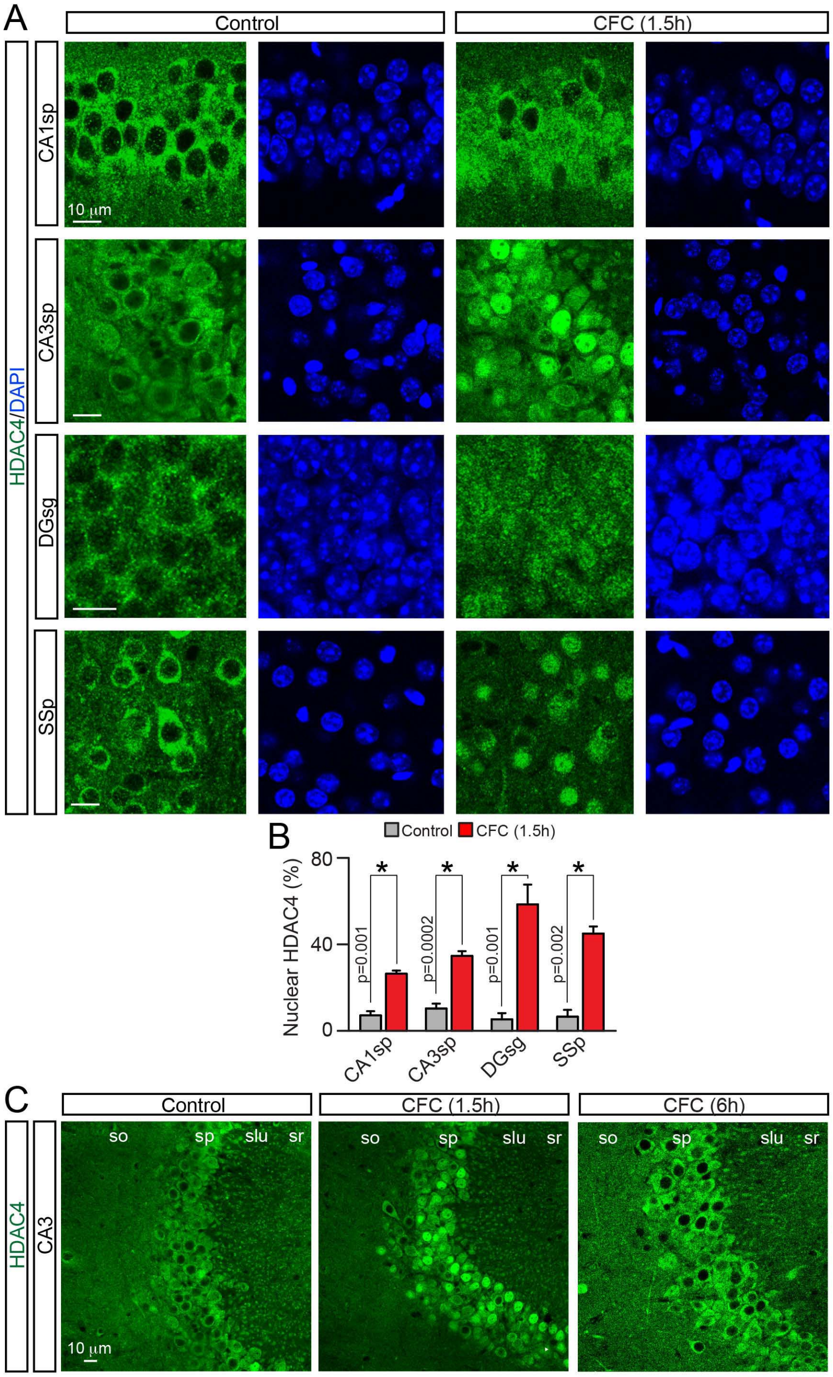


**Figure S1. Subcellular localization of native HDAC4 in brains of normal and fear-conditioned mice.** Brain sections were labeled with the antibody against HDAC4 and a nuclear marker, DAPI. **(A)** Typical high magnification confocal images of CA1sp, CA3sp, DGsg and SSp in samples from normal and foot-shocked animals (1.5 hours post-CFC). **(B)** Averaged percentages of cells with nuclear HDAC4 in indicated areas. Total cell numbers were quantified with DAPI. *n* = 3-4 mice/group. **(C)** Images of the CA3 demonstrate than nuclear accumulation of HDAC4 during associative learning is reversible.

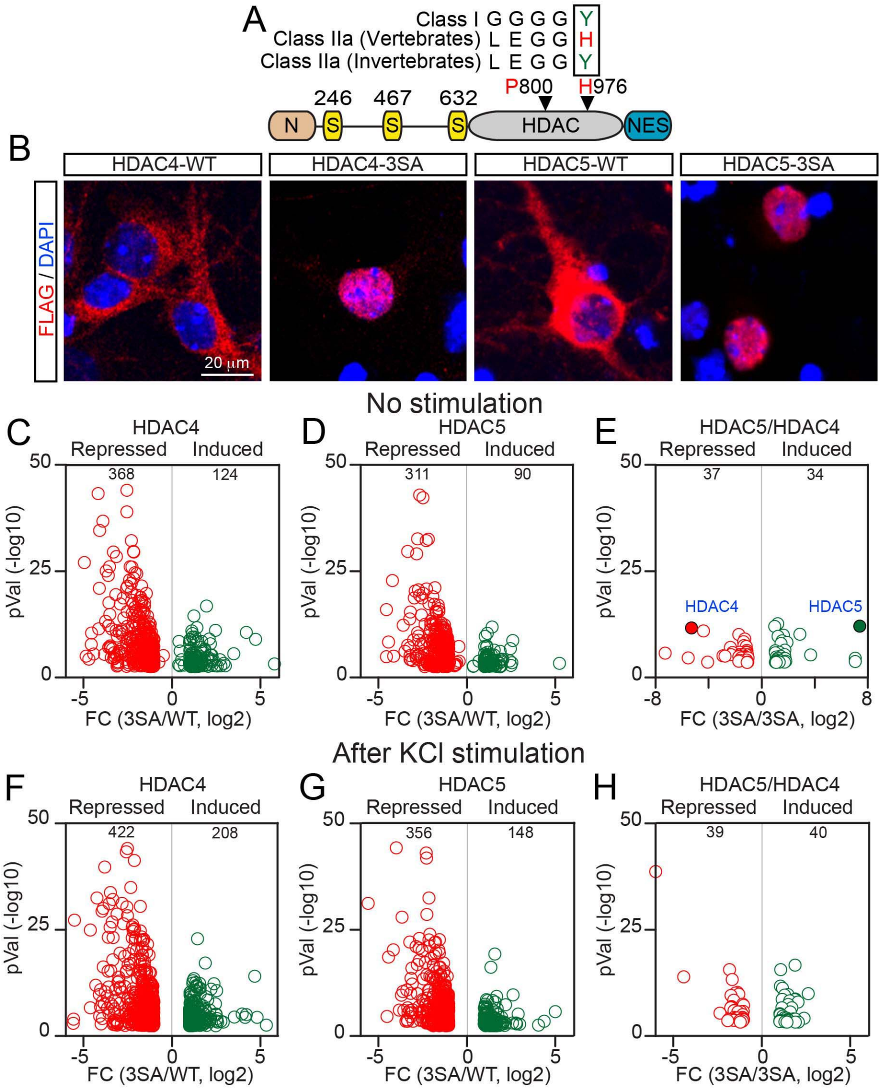


**Figure S2. (A)** Domain structure of HDAC4 (based on human sequence). The diagram depicts the N-terminal region, Serine residues 246, 467 and 632 whose phosphorylation promotes nuclear export, histone deacetylase domain (HDAC), nuclear export signal (NES), a frame shift mutation associated with mental retardation in humans (P800), and a mutation in the active site that diminishes catalytic activity of ε-N-acetyl lysine substrates (H976). A sequence alignment of these coding regions of conventional class I HDACs, and invertebrate and vertebrate class IIa HDACs is also shown (see also Lahm et al., 2007, Williams et al., 2010 and Sando et al., 2012). **(B** to **H)** Analysis of nuclear repressor activity of HDAC4 and HDAC5 *in vitro*. FLAG-tagged wildtype class IIa HDACs or their constitutively nuclear phosphorylation-deficient mutants (3SA) were broadly expressed in cortical neurons in primary cultures from lentiviruses under the control of the Synapsin promoter. Cultures were infected at 2 days *in vitro* (DIV2) and analyzed at DIV7. **(B)** Localization of recombinant proteins. Confocal images of neurons that were labeled with DAPI and the antibody against FLAG are shown. **(C** to **H)** Gene expression levels were examined by deep sequencing (RNA-seq) in unstimulated (panels C to E) and KCl-depolarized neurons (panels F to H). Volcano plots of all differentially expressed transcripts from 2 independent RNA-seq experiments are shown (total numbers are indicated in each panel). In panels C, D, F and G, fold changes (FC) are represented as log2 of 3SA/WT ratio for each HDAC isoform. Negative and positive values reflect repression and upregulation by 3SA mutants, respectively. Panels E and H demonstrate that only few genes are differentially expressed in neurons carrying HDAC4-3SA and HDAC5-3SA.

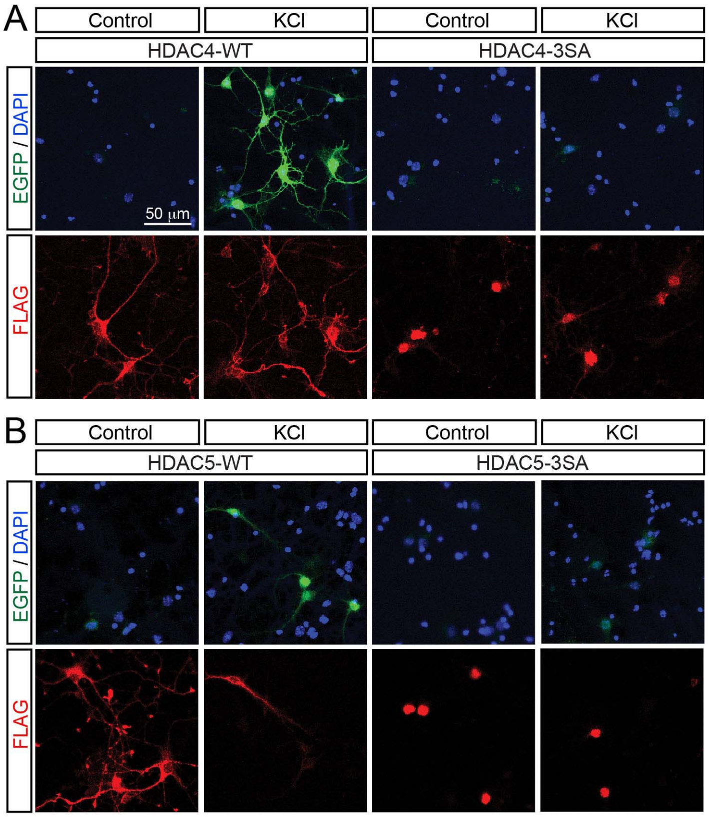


**Figure S3. Nuclear class IIa HDACs repress an activity-dependent reporter, E-SARE:EGFP. (A)** Confocal images of unstimulated and KCl-depolarized neurons that co-expressed E-SARE:EGFP with either cytoplasmic wildtype of nuclear gain-of-function 3SA HDAC4 construct. **(B)** Confocal images of unstimulated and KCl-depolarized neurons that co-expressed E-SARE:EGFP with either wildtype of nuclear gain-of-function form of HDAC5. In both panels, cultures were labeled with an antibody against FLAG and DAPI.

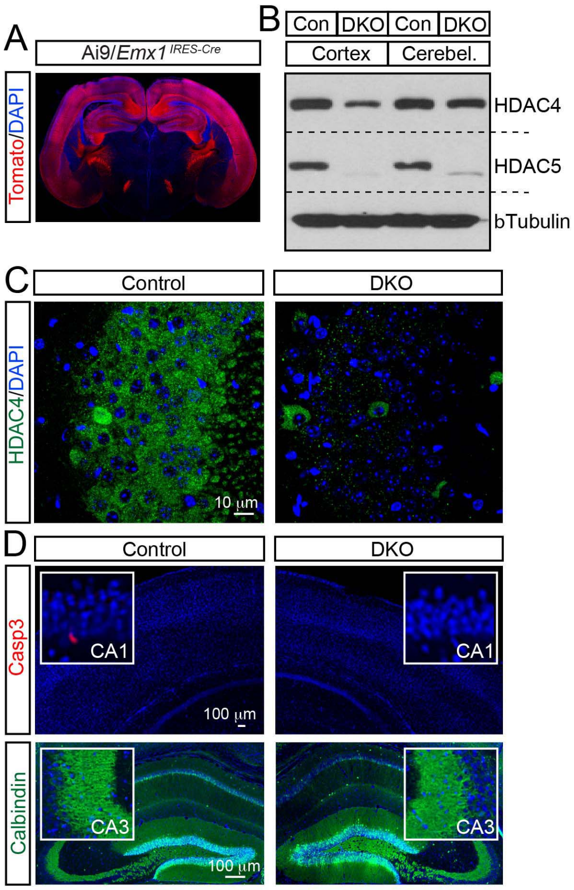


**Figure S4. (A)** Expression of tdTomato Cre reporter in *Ai9/Emx1*$^{IRES\text{-}Cre}$ mice. For detailed analysis, see Sando et al., Neuron, 2017. **(B)** Total protein homogenates from cortices and cerebella of adult wildtype (Control) and *Hdac4*$^{flox/flox}$/*Hdac5*$^{-/-}$/*Emx1*$^{IRES\text{-}Cre}$ (DKO) mice were probed by immunoblotting with antibodies against HDAC4, HDAC5 and βTubulin (as a loading control). Note that remaining expression of HDAC4 in DKO forebrain reflects lack of recombination in inhibitory GABAergic interneurons. **(C** and **D)** Brain sections from p60 control and DKO mice were labeled with DAPI and antibodies against HDAC4 (panel C) or cleaved Caspase 3 and Calbindin (panel D). Representative confocal images are shown.

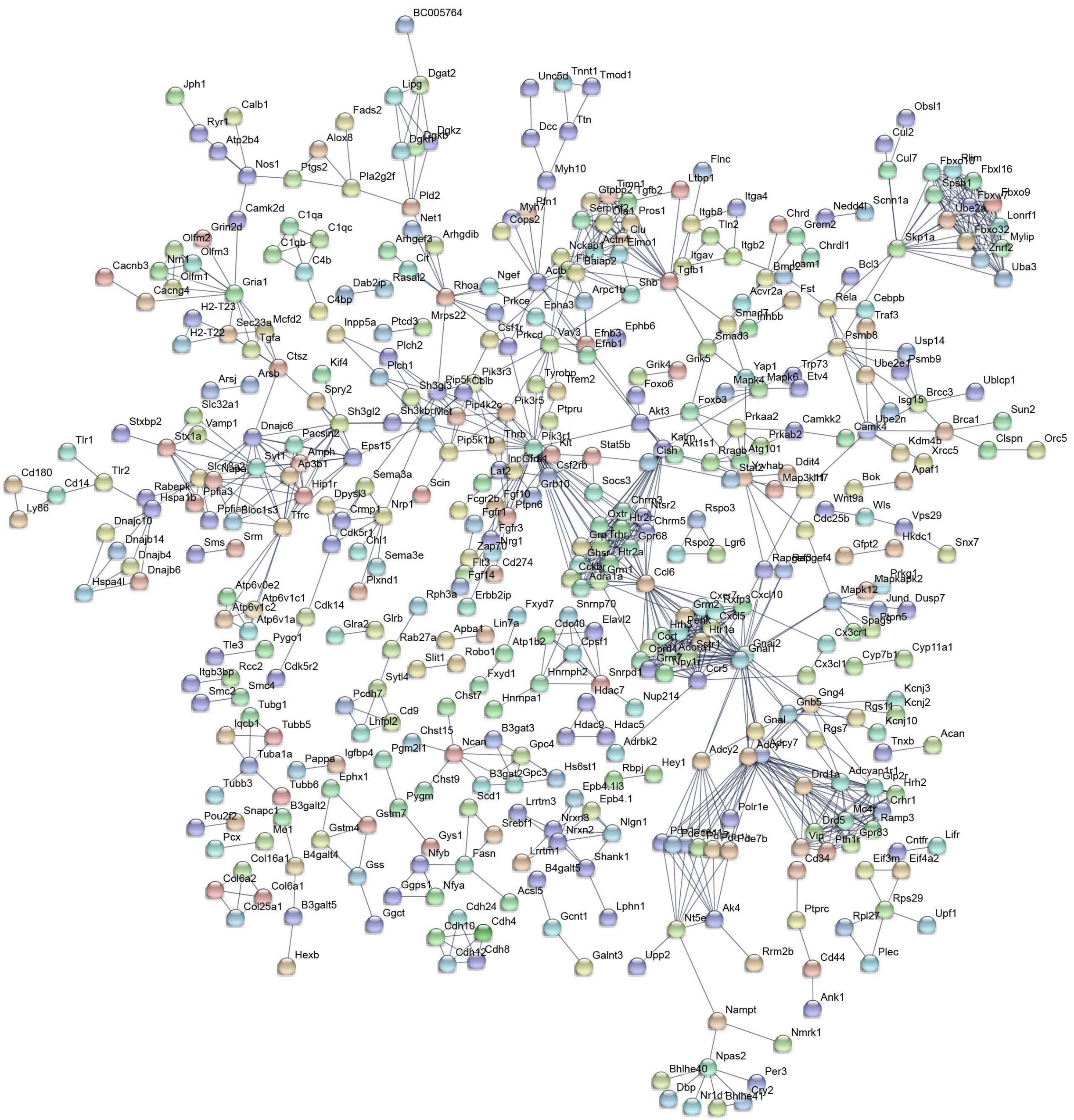


**Figure S5. STRING pathway analysis of differentially expressed genes in the hippocampus of DKO mice.** The network was generated at https://string-db.org using the highest confidence interaction score of 0.9. Disconnected nodes are not displayed.

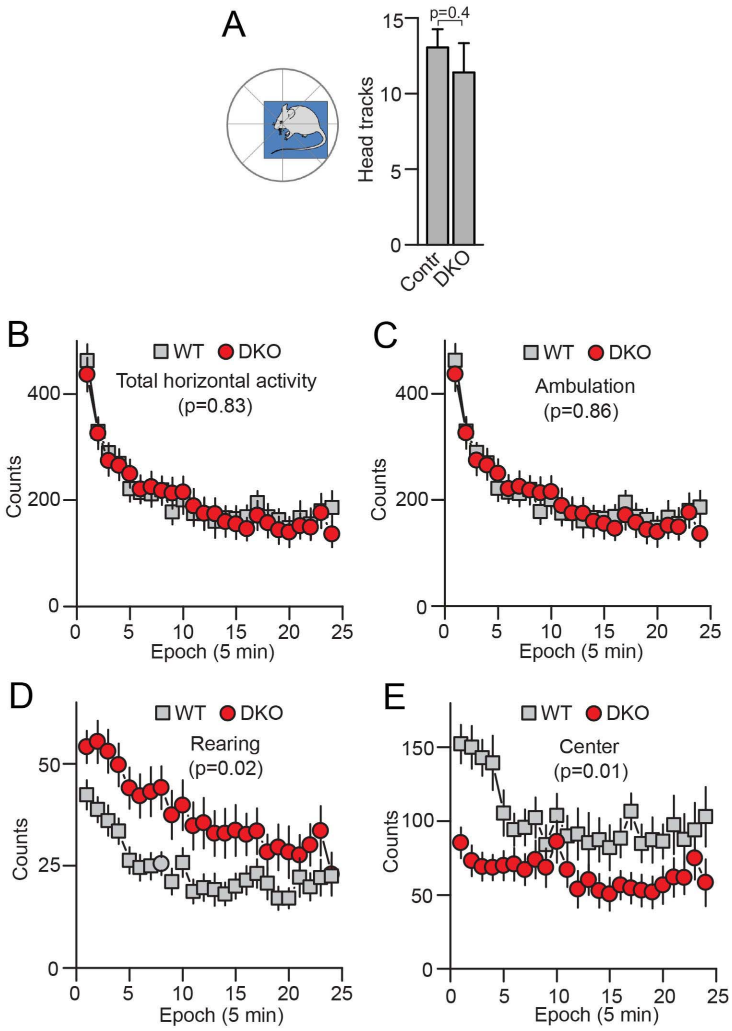


**Figure S6. (A)** Analysis of vision of p60 control and DKO mice. Schematic representation of the optomotor setup and averaged head tracks are shown. Control: $n = 19$; DKO: $n = 18$. **(B** to **E)** Detailed analysis of behavior in the open field. Quantifications of total horizontal activity (panel B), ambulation (panel C), rearing (panel D) and center activity (panel E) are shown. Control: $n = 19$; DKO: $n = 18$. All graphs are plotted as Mean ± S.E.M. $p$ values were determined by t-test.

A

MISLIAALAVDYVIGMENAMPWNLPADLAWFKRNTLNKPVIMGRYTWESIGRPLPGRKNIILSSQPSTDDRVTWV
KSVDEAIAACGDVPEIMVIGGGRVIEQFLPKAQKLYLTHIDAEVEGDTHFPDYEPDDWESVFSEFHDADAQNSHSY
CFEILERRGAPMSSQSHPDGLSGRDQPVELLNPARVNHMPSTVDVATALPLQVAPSAVPMDLRLDHQFSLPVAEP
ALREQQLQQELLALKQKQQIQRQILIAEFQRQHEQLSRQHEAQLHEHIKQQQEMLAMKHQQELLEHQRKLERHR
QEQELEKQHREQKLQQLKNKEKGKESAVASTEVKMKLQEFVLNKKKALAHRNLNHCISSDPRYWYGKTQHSSLDQ
SSPPQSGVSTSYNHPVLGMYDAKDDFPLRKTAAEPNLKLRSRLKQKVAERRSSPLLRRKDGPVVTALKKRPLDVTDS
ACSSAPGSGPSSPNNSSGSVSAENGIAPAVPSIPAETSLAHRLVAREGSAAPLPLYTSPSLPNITLGLPATGPSAGTAG
QQDAERLTLPALQQRLSLFPGTHLTPYLSTSPLERDGGAAHSPLLQHMVLLEQPPAQAPLVTGLGALPLHAQSLVG
ADRVSPSIHKLRQHRPLGRTQAAPLPQNAQALQHLVIQQQHQQFLEKHKQQFQQQQLQMNKIIPKPSEPARQPE
SHPEETEEELREHQALLDEPYLDRLPGQKEAHAQAGVQVKQEPIESDEEEAEPPREVEPGQRQPSEQELLFRQQAL
LLEQQRIHQLRNYQASMEAAGIPVSFGGHRPLSRAQASPASATFPVSVQEPPTKPRFTTGLVYDTLMLKHQCTCGS
SSSHPEHAGRIQSIWSRLQETGLRGKCECIRGRKATLEELQTVHSEAHTLLYGTNPLNRQKLDSKKLLGSLASVFVRL
PCGGVGVDSDTIWNEVHSAGAARLAVGCVVELVFKVATGELKNGFAVVRPPGHHAEESTPMGFCYFNSVAVAAK
LLQQRLSVSKILIVDWDVHHGNGTQQAFYSDPSVLYMSLHRYDDGNFFPGSGAPDEVGTGPGVGFNVNMAFTG
GLDPPMGDAEYLAAFRTVVMPIASEFAPDVVLVSSGFDAVEGHPTPLGGYNLSARCFGYLTKQLMGLAGGRIVLAL
EGGHDLTAICDASEACVSALLGNELDPLPEKVLQQRPNANAVRSMEKVMEIHSKYWRCLQRTTSTAGRSLIEAQTC
ENEEAETVTAMASLSVGVKPAEKRPDEEPMEEEPPLEFGGDYKDDDDK*

Blue: ecDHFR (DD)
Black: HDAC4 (S/A mutations are shown in red)
Yellow: FLAG tag

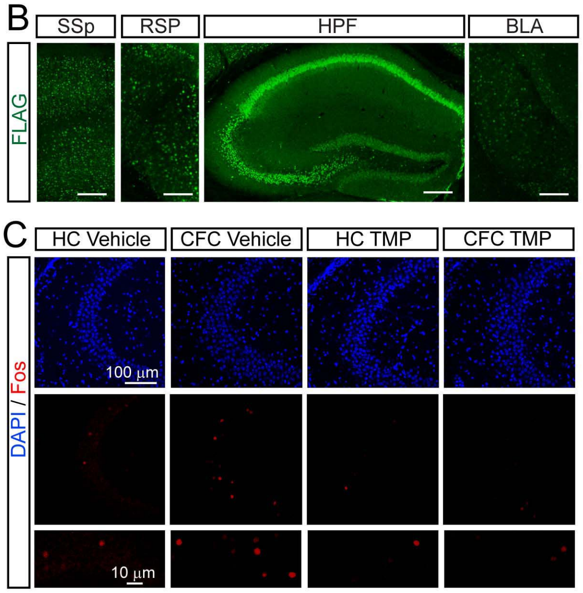


**Figure S7. Characterization of *R26$^{floxStop\text{-}DD\text{-}nHDAC4}$/Camk2α$^{Cre}$* mice. (A)** Primary aminoacid sequence of DD-nHDAC4 fusion protein. **(B)** Immunofluorescent imaging analysis of expression of DD-nHDAC4-F in different brain regions. SSp: Primary somato-sensory cortex; RSP: Retrosplenial cortex; HPF: Hippocampus; BLA: Basolateral amygdala. **(C)** Acutely stabilized DD-nHDAC4 represses ERGs. *R26$^{floxStop\text{-}DD\text{-}nHDAC4}$/Camk2α$^{Cre}$* mice were given single doses of vehicle or TMP. Injected animals were either maintained in home cages (HC) or subjected to associative learning via CFC (3 hours post-injection). 1.5 hours later, brains were isolated, sectioned, and imaged after labeling with DAPI and an antibody against Fos. Typical images of the CA3 are shown.

**Table S1** (Separate Excel file).

RNA-seq analysis of gene expression in cultured neurons.

**Table S2** (Separate Excel file).

RNA-seq analysis of gene expression in hippocampi of control and DKO mice.
